# Genetic dissection of transcription start site selection by RNA Polymerase II in *Saccharomyces cerevisiae*

**DOI:** 10.1101/2025.10.24.684399

**Authors:** Payal Arora, Kieran F. Brennan, Marcie H. Warner, Craig D. Kaplan

**Author notes:** Address for correspondence: Department of Biological Sciences, University of Pittsburgh, Pittsburgh, PA 15260, Life Science Annex 202, Fifth and Ruskin Avenues, Pittsburgh, PA 15260.

## Abstract

Transcription initiation by RNA Pol II is driven by a preinitiation complex (PIC) comprising Pol II and general transcription factors (GTFs): TFIIA, TFIIB, TFIID, TFIIE, TFIIF and TFIIH. In *Saccharomyces cerevisiae*, transcription start site (TSS) selection proceeds by a promoter scanning mechanism where the PIC scans downstream for TSSs. To determine which factors participate in TSS selection by promoter scanning, we designed and implemented forward genetic selections using initiation-sensitive genetic reporters. These reporters were of two types, either being sensitive to upstream or downstream TSS shifts, allowing detection of alleles that affect promoter scanning in different directions. From >1000 candidates, we identified three primary classes of mutants: existing and novel mutants within known PIC components, including novel alleles of multiple TFIIH subunits; mutants that alter promoter scanning indirectly through cellular conditions (GTP levels and Mn^2+^ levels); and mutants in chromatin and transcription elongation-related factors. Genome-wide analysis using TSS mapping shows that tested PIC mutants alter TSS usage globally across promoters while tested chromatin and transcription elongation mutants had much more limited effects. These studies expand the spectrum of mutants able to perturb initiation by promoter scanning and suggest potential plasticity in the TFIIH structure that may be part of the evolution of this initiation mechanism.

## INTRODUCTION

Transcription initiation in eukaryotes requires assembly of a large pre-initiation complex (PIC) comprising RNA Polymerase II (Pol II) and general transcription factors (GTF) (CONAWAY AND CONAWAY 1993; ORPHANIDES *et al*. 1996; ROEDER 1996; KORNBERG 2007). Together these components, in conjunction with transcriptional activators and coactivators effect promoter DNA opening, identification of the transcription start site (TSS), and de novo initiation of RNA synthesis using DNA-templated NTP substrates (KAPLAN 2013; ROEDER 2019). Additionally, a large number of other factors are recruited to promoters, including chromatin remodelers and factors functioning in the transition from initiation to elongation (RANDO AND WINSTON 2012). During PIC assembly, TBP and TFIIB bind to the core promoter transiently with TFIIA stabilizing TBP-DNA interactions. Pol II is then recruited to core promoter along with GTFs TFIIF and TFIIE stabilizing the TFIIB-TBP-DNA interactions. Thereafter, TFIIH is recruited into the PIC to drive open complex formation (promoter melting) and subsequent transcription initiation (BAEK *et al*. 2021).

In eukaryotes, the majority of promoters specifies multiple TSSs. In most eukaryotes, a TSS is stereotypically positioned approximately 30-31 basepairs downstream from a site of TBP binding (a canonical TATA-box or otherwise AT-rich sequence) (LIFTON *et al*. 1978; BREATHNACH AND CHAMBON 1981; DUDNYK *et al*. 2024). It is proposed that each TSS is supported by an associated PIC assembly point upstream (KADONAGA 1990; SMALE AND KADONAGA 2003; SANDELIN *et al*. 2007; KADONAGA 2012; VO NGOC *et al*. 2017a; VO NGOC *et al*. 2017b; LUSE *et al*. 2020; DUDNYK *et al*. 2024). In contrast, transcription initiation in the budding yeast *Saccharomyces cerevisiae* occurs through a promoter scanning mechanism (GIARDINA AND LIS 1993; KUEHNER AND BROW 2006; FAZAL *et al*. 2015; QIU *et al*. 2020). The driving factors for this novel initiation mechanism can only be speculated upon but given the loss of most splicing in budding yeast and its highly compact genome, scanning may allow for 5′ UTR diversification that cannot be accomplished by alternative splicing coupled to alternative promoter usage. Intriguingly, PIC components and their assembled structure are highly conserved between yeast and humans (BURATOWSKI *et al*. 1988; HE *et al*. 2016; AIBARA *et al*. 2021; SCHILBACH *et al*. 2021; TOMKO *et al*. 2021). While DNA melting by the yeast PIC occurs proximal to its site of binding as in other eukaryotes (GIARDINA AND LIS 1993), the yeast PIC scans downstream for TSSs, which are generally located 40-120 bp or further downstream of the TATA box, when present (STRUHL 1987; PARK *et al*. 2014; LU AND LIN 2019; LU AND LIN 2021). TFIIH, required for Pol II promoter DNA opening in all eukaryotes, is proposed to use its DNA translocase activity to continue pumping downstream DNA into the Pol II active site to enable promoter scanning and usage of downstream TSSs (HAMPSEY 2006; MILLER AND HAHN 2006; FAZAL *et al*. 2015; FISHBURN *et al*. 2015).

Promoter scanning identifies TSSs based on their inherent sequence characteristics together with the contributions of initiation factor and Pol II activities. Yeast share a universal eukaryotic preference for Y_-1_R_+1_ as TSSs initiating at a purine (R, representing A or G), have a pyrimidine (Y, representing C or T) just upstream of the TSS, and with an additional species-specific contribution of the -8 position and of additional bases (HUROWITZ AND BROWN 2003; ZHANG AND DIETRICH 2005; MIURA *et al*. 2006; NAGALAKSHMI *et al*. 2008; ARRIBERE AND GILBERT 2013; PELECHANO *et al*. 2013; MALABAT *et al*. 2015; LU AND LIN 2019; QIU *et al*. 2020; LU AND LIN 2021; ZHU *et al*. 2024). Scanning in yeast was conceptualized by Kuehner and Brow (KUEHNER AND BROW 2006) where TSSs have innate efficiencies of usage, and the alteration of initiation factor activity can alter individual TSS usage and the distribution of TSS usage at promoters (KUEHNER AND BROW 2006; QIU *et al*. 2020; ZHAO *et al*. 2021). Genetic and molecular experiments have shown that TSS selection in yeast is sensitive to mutation in multiple PIC components (*e.g.* Pol II and GTFs (PINTO *et al*. 1992a; PINTO *et al*. 1994; FAITAR *et al*. 2001; GHAZY *et al*. 2004; FREIRE-PICOS *et al*. 2005; KHAPERSKYY *et al*. 2008; JIN AND KAPLAN 2014; QIU *et al*. 2020; ZHAO *et al*. 2021) and PIC-associated components (*e.g.* the conserved factor Sub1, which is the yeast homolog of PC4 (KNAUS *et al*. 1996; WU *et al*. 1999; SIKORSKI *et al*. 2011; BRABERG *et al*. 2013). Mutations in the active site of Pol II that increase Pol II catalytic activity in vitro increase efficiency of initiation at each available TSS (KAPLAN *et al*. 2012; JIN AND KAPLAN 2014; QIU *et al*. 2020). With each TSS being more efficient in promoting initiation, the flux of scanning Pol II can be exhausted earlier than WT, leading to an observed upstream shift in observed TSS usage (KUEHNER AND BROW 2006; QIU *et al*. 2020). Similarly, decrease in Pol II catalytic activity causes reduced efficiency of individual TSSs, resulting in increases in the flux of scanning Pol II reaching downstream positions, and an observed downstream shift in TSS usage (KUEHNER AND BROW 2006; QIU *et al*. 2020).

In contrast to Pol II, because TFIIH is hypothesized to drive scanning through its Ssl2 translocase subunit (FAZAL *et al*. 2015; FISHBURN *et al*. 2015; TOMKO *et al*. 2017; TOMKO *et al*. 2021; ZHAO *et al*. 2021), the effects of mutations in Ssl2 on TSS distributions have been interpreted as being due to altered TFIIH processivity (ZHAO *et al*. 2021). *ssl2* alleles that confer downstream shifts in TSS distributions have been interpreted as increasing scanning processivity due to a higher probability of scanning reaching downstream positions. In contrast, *ssl2* alleles that confer upstream shifts in TSS distributions appear to do so by limiting scanning processivity, meaning that downstream positions are reached at a lower probability (KUEHNER AND BROW 2006; ZHAO *et al*. 2021). While TFIIH and Pol II are the primary activities driving TSS selection, how GTFs contribute to either efficiency or processivity pathways or through additional mechanisms is not known (JIN AND KAPLAN 2014; QIU *et al*. 2020).

Mutations in other GTFs like TFIIB (PINTO *et al*. 1992a; PINTO *et al*. 1994; BANGUR *et al*. 1997; PARDEE *et al*. 1998; FAITAR *et al*. 2001; JIN AND KAPLAN 2014) and TFIIF (SUN AND HAMPSEY 1996; GHAZY *et al*. 2004; JIN AND KAPLAN 2014) also affect TSS distributions, and among those tested, to do so genome wide (QIU *et al*. 2020; BASNET *et al*. 2026), as do tested Pol II and *ssl2* alleles of TFIIH (QIU *et al*. 2020; ZHAO *et al*. 2021). Mutations in TFIIB cause downstream shifts to TSS usage in ways that appear consistent with both TFIIB and the Pol II active site determining initiation efficiency at all TSSs (WU *et al*. 1999; FAITAR *et al*. 2001; KOSTREWA *et al*. 2009; JIN AND KAPLAN 2014; QIU *et al*. 2020). Additionally, published mutations in TFIIF, the GTF recruiting Pol II to the core promoter, have also been shown to shift TSSs upstream (SUN AND HAMPSEY 1995; GHAZY *et al*. 2004; FREIRE-PICOS *et al*. 2005; JIN AND KAPLAN 2014; QIU *et al*. 2020). Because specific alleles of TFIIF and TFIIB have been found to mutually suppress each other (GHAZY *et al*. 2004; FREIRE-PICOS *et al*. 2005) and also additively interact with Pol II alleles (JIN AND KAPLAN 2014; QIU *et al*. 2020), we have proposed that at least some TFIIF mutants affect TSS selection by functioning alongside the Pol II active site in promoting initiation efficiency (GHAZY *et al*. 2004; FREIRE-PICOS *et al*. 2005; JIN AND KAPLAN 2014). The interface between Pol II and TFIIH is also important for scanning and has been proposed to be a key factor in the evolution of the scanning mechanism (YANG *et al*. 2022; YANG *et al*. 2024; BASNET *et al*. 2026). TFIIH is tethered to the PIC via the RING finger domain of Tfb3 (TFIIK kinase subdomain of TFIIH) through interactions with TFIIE and the Rpb7 subunit of Pol II (SCHILBACH *et al*. 2017; YANG *et al*. 2022; YANG *et al*. 2024). In yeast, mutations in both Rpb7 (BRABERG *et al*. 2013) and Tfb3 (YANG *et al*. 2024) can confer upstream shifts to TSS usage *in vivo*, consistent with a defective scanning process (BASNET *et al*. 2026). Factors outside of PIC such as chromatin and elongation factors have also been implicated to affect promoter choice and potentially TSS selection.

PIC factors (Pol II and GTFs) and PIC-associated factors like Sub1 have been genetically identified as contributing to TSS-selection during promoter scanning. What is not known is how many additional factors might also contribute to the process of TSS selection. Are there additional factors that cooperate or contribute to TSS selection but have not been structurally located in the PIC akin to Sub1? What are the surfaces and networks within the PIC that contribute to TSS selection by promoter scanning? Finally, promoter chromatin structure can easily be imagined to affect PIC function and TSS distribution profiles because most yeast promoters have TSSs that map within boundaries of the +1 nucleosome (RANDO AND WINSTON 2012). We have aimed to take an unbiased approach to identify any additional factors that participate in TSS selection during initiation by promoter scanning using forward genetic selections (BROW 2019; WANG *et al*. 2025). Through the use of existing (BERROTERAN *et al*. 1994) and novel genetic reporters, we have identified ∼300 mutations in proteins within and outside the PIC affecting TSS selection, including a number of alleles of Pol II elongation/chromatin genes, *eg. SPT2, SPT4, SPT5, SPT6, SPT16, SPN1, BUR2*, and histones. We then examined a subset of the novel alleles in genetic interaction studies to probe relationships between mutant alleles. Finally, we examined the genome-wide effects of selected PIC and chromatin/elongation-class alleles for effects on TSS-selection. We have found that tested GTF alleles alter TSS usage genome wide as predicted from effects on promoter scanning in general. In contrast, chromatin or Pol II elongation-class alleles did not show similar widespread effects. We interpret this class of mutant as acting on TSSs in a promoter-selective fashion.

## MATERIALS AND METHODS

### Yeast strains and oligos

Yeast strains and oligos used in this study are listed in **Table S2**.

### Yeast media

Yeast media used in the current studies have been prepared as previously described in (AMBERG *et al*. 2005; KAPLAN *et al*. 2012; JIN AND KAPLAN 2014; MALIK *et al*. 2017; ZHAO *et al*. 2021). Briefly, all solid media contain 2% w/v agar (BD); YPD is 1% w/v yeast extract (Gibco™ Bacto™), 2% w/v peptone (Gibco™ Bacto™) and 2% w/v glucose (VWR) supplemented with 0.15 mM adenine (Sigma) and 0.4 mM tryptophan (Sigma) for solid medium (not supplemented when liquid); YPD+3% formamide contains 3% v/v formamide (Ambion); YP Raffinose (YPRaf) and YP Raffinose Galactose (YPRafGal) contain 2% w/v raffinose (Amresco) (YPRaf) or 1% w/v galactose (Sigma) and 2% w/v raffinose (YPRafGal). Both YPRaf and YPRafGal are additionally supplemented with 1 µg/ml of (Sigma) Antimycin A to prevent aerobic respiration. Synthetic minimal media (SC-) were what is referred to as “Hopkins mix” with minor alterations as previously noted (AMBERG *et al*. 2005; KAPLAN *et al*. 2012) (amino acid dropout mix, 0.17% w/v yeast nitrogen base w/o amino acids and 0.5% w/v ammonium sulphate (BD), 2% glucose), with the appropriate dropouts. Mycophenolic acid (MPA) (ThermoScientific chemicals) containing SC media were supplemented at 20 µg/ml. Other drugs in the respective media were: G418 (GoldBio) and clonNAT (GoldBio) at 200 µg/ml, hygromycin (GoldBio) at 300 µg/ml and canavanine (Sigma) at 60 µg/ml. Synthetic minimal media containing G418, clonNAT or hygromycin are prepared as above with 0.1% w/v of monosodium glutamate (MSG) replacing ammonium sulphate. Drug 3-aminotriazole (3AT) (Sigma) a competitive inhibitor of His3p was used in some SC-His media at varying concentrations (later noted), for setting the threshold for a visible His^+^ phenotype.

### Genetic selections

Each of the genetic selection strains possessing initiation reporters was grown overnight in liquid YPD. Cultures were then washed twice in sterile distilled water to remove residual amino acids before plating on respective selection media. Selection media for the reporters were SC-His + 2mM 3AT for *imd2Δ::HIS3* and SC-His + 0.1mM 3AT for *cyc1-1019::HIS3*, *adhΔ50::HIS3* and *hmo1Δ38::HIS3*. For dual selection of His^+^ Lys^+^ candidates for downstream reporters, conditions used were SC-His-Lys +5mM 3AT for *imd2Δ::HIS3* SC-His-Lys +0.1mM 3AT, Cells were spread using sterile 3mm glass beads at 10^8^ cells per plate. Multiple colonies of each parent strain were inoculated for genetic selection to increase chances of independent candidates. Plates were then kept in a lightly sealed plastic bag to prevent drying out and incubated at 30°C for ∼7 days until suppressor colonies appeared.

### Single spot phenotyping

Suppressor candidates were picked from selection plates and spotted onto solid YPD petri plate in 8x6 grid (48-well format) containing 2 parents and 46 candidates apiece for the purpose of candidate rescreening and additional phenotyping. Briefly, after growth on YPD, cells were transferred into 100 µl of sterile distilled water in a 96-well tray using a 48-pin replicator, followed by a second dilution of 6 µl of this cell suspension into 150µl of water. A sterilized 48-pin replicator was then used to spot from the 150µl cell suspension onto phenotyping media. Phenotype scoring timing was day 2 for YPD 30°C and YPD 37°C, day 3 for YPD+3% formamide and synthetic complete medium (SC), day 4 for YPRaf, day 5 for SC-Lys and YPD at 16°C and day 6 for YPRafGal. Reporter phenotypes on SC-His +/- 3AT were scored on days 3 through 5 depending on strength and SC-His-Lys +/- 3AT on day 5. **Scoring:** Scoring was performed as in (JIN AND KAPLAN 2014). Single phenotype spots were given scores from 0 to 5, where 0 means no growth and 5 is WT parent growth on complete medium. For conditional phenotypes where growth is indicative of mutant phenotype (i.e. Spt^-^ phenotype or His^+^ phenotypes in reporter strains), scoring was 0 for no growth (WT) with positive numbers indicating increasing growth reflecting strength of mutant phenotype. Relative scores of mutants were first calculated per condition by Score_mutant_-Score_WT_ (Relative score_mutant: condition_). These relative scores were then corrected to respective control growth condition for estimating mutant score corrected to control condition (e.g.: Relative score_mutant: YPRafGal_ - Relative score_mutant: YPRaf_). These would assign positive values to mutants where growth is better than WT, negative to those where growth is worse in an any condition and a 0 to WT.

### Mutation validation by re-introduction

Mutations were reintroduced into a clean parental strain possessing relevant initiation reporter to validate effects of candidate mutations on reporter expression. Mutations were amplified from suppressor strains and fused with a drug resistance marker by PCR sewing to generate a *mutation::drug resistance marker* cassette and reintroduced into parent strains (JANKE *et al*. 2004). The marker used was *kanMX6* containing an insulated resistance cassette (pFA6a-KanMX6-ins2, Addgene 195039) to minimize potential effects of marker expression on adjacent loci (POWERS *et al*. 2022). Multiple independent transformants were phenotyped and genotyped to assess correlation of reporter suppression and introduction of the mutant genotype along with the resistance marker. The marker was also used to tag mutation loci to enable synthetic genetic array analysis and selection of mutant loci through genetic crosses. CRISPR based mutagenesis was performed to generate *rpb10* and *tfg1* mutations in the genome as described in (LAUGHERY *et al*. 2015).

### Diploid generation

Complementation assays and genetic interaction studies were performed with diploid generation. Diploids were generated by mating suppressor candidates against parent or mutant strains of opposite mating type, followed by double selection of individual haploid markers for selecting diploids. Briefly, candidate suppressors, pre-integrated with auxotrophic markers (*LEU2* for *MATalpha* and *TRP1* for *MATa*) were mated with the opposite mating type of the required strains with a second auxotrophic selection marker; or appropriate drug resistance markers were used for diploid selection for genetic interactions. Mating was performed by transferring solid YPD-grown candidates to pre-aliquoted 100 µl-per-well of saturated culture of test strain per well in a 96-well plate using a 48-pin (8x6) replicator and spotting onto solid YPD using the same replicator. Post incubation for 2 days at 30°C on plates, cells were transferred into 100µl of pre-aliquoted sterile distilled water with the 48-pin replicator and resuspended by shaking of replicator followed by transfer onto a double-selection media. These were incubated at 30°C to select for diploids for 2 days.

### Double mutant generation by synthetic genetic array

Double mutants were generated from diploids of “tagged” single mutants mated in high throughput following protocol from the Boone lab (https://boonelab.ccbr.utoronto.ca/pdf/SGA_protocol_final.pdf). Briefly, a panel of *MAT⍺ single mutation 1::drug resistance marker 1* strains was mated to a query *mutation 2::drug resistance marker 2 MATa* strain possessing a *MATa* marker *can1*Δ*::STE2pr-URA3,* and diploids were selected by double drug resistance as mentioned. After two days of incubation on double drug diploids were transferred from solid medium directly onto solid enriched sporulation media (1% potassium acetate enriched with auxotrophic amino acids in 2% agar) for 5 days at 27°C. After sporulation, sporulated spots were transferred into 50 µl sterile distilled water in 96 well plate using a pinner, resuspended and spotted onto haploid selection media (SC-Ura-Arg+MSG+canavanine), single mutant selection media (SC-Ura-Arg+MSG+canavanine + drug 1 or drug 2) and double mutant (SC-Ura-Arg+MSG+canavanine+drug 1+drug 2) selection media in parallel instead of the traditional approach where mutants (single or double) are selected after *MATa* meiotic progeny selection in tandem. We have observed that sicker haploids are lost if selections are performed in tandem.

### Genomic DNA sequencing

Candidate mutations were identified via AmpliSeq-based custom amplicon sequencing or whole genome sequencing (WANG *et al*. 2025). Genomic DNA was isolated from candidates using PureLink™ Pro 96 Genomic DNA Purification Kit and YeaStar genomic DNA kit from Zymo research for AmpliSeq based sequencing. The Plasmid miniprep kit from Zymo research, customized with bead lysis was used for genomic DNA isolation for whole genome sequencing. AmpliSeq libraries were prepared and sequenced by the Texas A&M TIGGS genomics core facility. AmpliSeq libraries were prepared following AmpliSeq for Illumina protocol for a custom oligo pool comprising 1917 encompassing 310 genes. These genes comprise ORFs of transcription machinery and transcription-associated genes. Libraries were sequenced in batches of 96 samples using full Illumina MiSeq of 15 million reads by 250bpx2 run to aim for an average coverage of ∼80X per amplicon. Whole genome sequencing of samples was performed by Seqcoast or Seqcenter. Each genomic DNA sample was sequenced at a depth of 400Mbp yielding an average coverage of ∼30X.

### AmpliSeq data processing

Paired-end sequencing fastq files were first trimmed for adaptors at the 3′ end, followed by 3′ and 5′ oligo trimming (amplification oligos do not contribute towards mutation detection) by cutadapt v2.1 (MARTIN 2011). These were then processed by MutantHuntWGS v1.1 (ELLISON *et al*. 2020), a container-based package deploying predefined tools for alignments and variant calling. Briefly, oligo and adaptor-free paired-end fastq files were aligned to *Saccharomyces cerevisiae* S288C reference genome R64-2-1 by bowtie2 v2.2.9 (LANGMEAD AND SALZBERG 2012) resulting in bam alignment files followed by bam sorting and indexing by SAMtools v1.3.1 (LI *et al*. 2009; DANECEK *et al*. 2021). Genotype likelihoods were calculated followed by variant calling by BCFtools v1.3.1 (DANECEK *et al*. 2021) using haploid ploidy file provided by MutantHuntWGS v1.1 (ELLISON *et al*. 2020) (ELLISON *et al*. 2020). This resulted in vcf files for all sequenced candidates. Mutant variants in mutant vcf files were differentially called relative to parent vcf files by VCFtools v 0.1.14 (DANECEK *et al*. 2011). Resulting differential mutants were annotated using SNPeff v4.3p (CINGOLANI *et al*. 2012) and SIFT4G (VASER *et al*. 2016), which uses EF4.74 *Saccharomyces cerevisiae* library for annotating severity of amino acid change. For WGS, fastq reads were analyzed by Seqcenter and Seqcoast for mutation detection by breseq (DEATHERAGE AND BARRICK 2014). *Saccharomyces cerevisiae* S288C reference genome R64-1-1 was used for annotation which generated a list of variants. Variants specific to suppressor candidate were curated by a custom shell script comparing mutant variants to parent variants.

### STRIPE-seq

Total RNA was isolated from 50 mL of mid-log yeast culture with density of 1X10^7^ cells/mL as described previously (SCHMITT *et al*. 1990) and resuspended in 90 µL nuclease free water. RNA samples were in-column DNase I treated using the RNeasy mini kit from Qiagen with 100 µg RNA input per column. DNased RNAs (10 µg per sample) were depleted of rRNA using bioltinylated riboPOOL kit for *Saccharomyces cerevisiae* from siTOOLS Biotech, purified by Dynabeads MyOne Streptavidin C1 and eluted in 9 µL nuclease free water. STRIPE-seq libraries were then generated from rRNA depleted RNAs using 200 ng RNA per sample as previously published (POLICASTRO *et al*. 2020) with our modifications. Briefly, 200 ng of each RNA sample, split into two 100 ng reactions were terminator exonuclease (TEX) treated to remove uncapped RNA. Pooled, TEXed reactions were then reverse transcribed using a random 10-mer with partial homology to P7 adaptor RTO (GCCTTGGCACCCGAGAATTCCANNNNNNNNNN) instead of a complete P7:oligo-random pentamer (modification; P7 adaptor added during final library amplification). Non-templated cytosines at the 3′ end of cDNA then allow annealing of a template switching oligo, containing partial P5 adaptor, 9-base UMI (instead of 8-base UMI) and a linker (TATAGGG) (Biotin-GTTCAGAGTTCTACAGTCCGACGATCNNNNNNNNNTATArGrGrG). After template switching, libraries were amplified by dual indexing primers with complete P5 and P7 adpaters. Libraries (>200-800 bp) were extracted from 3% polyacrylamide TBE gel to remove oligos and then sequenced by Illumina NextSeq 2000 system with 1X50 bp sequencing cycle. Sequencing reads were delivered demultiplexed.

### STRIPE-seq analysis

STRIPE-seq fastq files were processed as TSS-seq reads we had previously published (ZHAO *et al*. 2021) with STAR alignment and additional bam processing followed previously published of goSTRIPES and TSRexploreR packages (POLICASTRO *et al*. 2020; POLICASTRO *et al*. 2021; POLICASTRO AND ZENTNER 2022). Fastq files were filtered for quality with fastx-toolkit v 0.0.14 (-q 20 -p 75) (http://hannonlab.cshl.edu/fastx_toolkit/), followed by trimming of adaptors by trimmomatic v 0.38 (BOLGER *et al*. 2014) and then 9 base UMI extraction by umi-tools v 1.0.0 (SMITH *et al*. 2017). Reads were then trimmed of linker TATAGGG and then trimmed to 25 bases by cutadapt v 2.10 (MARTIN 2011) before local alignment by STAR v 2.7.9a (DOBIN *et al*. 2013) generating bam files. Bam files with uniquely mapping reads were then filtered of unfavorably flagged reads by samtools v 1.9 (samtools view –F 3844) (LI *et al*. 2009; DANECEK *et al*. 2021) followed by deduplication by umi-tools v 1.0.0 (SMITH *et al*. 2017). Deduplicated and indexed bam files were then processed by the protocol followed by the Zentner lab (POLICASTRO *et al*. 2020; POLICASTRO *et al*. 2021; POLICASTRO AND ZENTNER 2022). Briefly, reads with more than 3 soft-clipped 5′ bases are listed and filtered out by picard v 2.18.12 (Picard Toolkit 2019) following GoSTRIPES (Robert Policastro). Indexed filtered bams were then processed by TSRexploreR (POLICASTRO *et al*. 2021) for G correction to generate bedgraphs for 5′ ends of reads (TSSs) that we processed to split into + and – strand bedgraphs and downstream analyses following our TSS-seq pipeline (ZHAO *et al*. 2021). Pearson correlation matrices were generated for bedgraphs of all libraries using our in-house Rscript. Promoter matrices representing TSSs were generated comprising 401 base window (250 bases upstream and 150 bases downstream from median TSS position including the median) across 5979 annotated promoters (RHEE AND PUGH 2012). These matrices were utilized for estimating median and spread (base distance between 10^th^ and 90^th^ percentile) of TSS distributions across the 5979 promoter and eventually Δmedian and Δspread of mutants relative to WT for differences (ZHAO *et al*. 2021). Promoter matrices were row normalized before generating average matrices across replicates and estimating average median and average spread values for calculating average Δmedian and Δspread values relative to average WT. Top 2829 promoters with expression cutoff of 10 were selected based on library normalized average of all WTs sorted by expression: sum of TSSs per 401 base promoter window.

## RESULTS

### TSS-shifting mutants identified by transcription initiation genetic reporters

To understand the genetic requirements for initiation by promoter scanning, we established genetic selection systems for the isolation of transcription start site-shifting mutants. We deployed four initiation reporter alleles designed or chosen because expression from the reporter is sensitive to altered TSS selection: *imd2Δ::HIS3, cyc1-1019::HIS3*, *adhΔ50::HIS3,* and *hmo1Δ38::HIS3* (**Fig. 1**). These reporters comprise native (*imd2Δ::HIS3*) (MALIK *et al*. 2017) or engineered (*cyc1-1019::HIS3*, *adhΔ50::HIS3* and *hmo1Δ38::HIS3*) (FURTER-GRAVES AND HALL 1990; PINTO *et al*. 1992a; PINTO *et al*. 1992b) promoters that have each been fused to the *HIS3* ORF. In wild type cells, they generally do not express sufficient levels of *HIS3* gene product to confer growth on medium lacking histidine (the His^-^ phenotype). In the presence of a TSS selection mutant conferring an upstream or downstream shift appropriate to the reporter configuration, a shift in TSS usage leads to expression of a functional *HIS3* mRNA, allowing cells to grow on medium lacking histidine (the His^+^ phenotype) (**Fig. 1a**). We tuned our phenotypic selections through the use of the drug 3-aminotriazole (3AT), a competitive inhibitor of His3p (KLOPOTOWSKI AND WIATER 1965; STRUHL AND DAVIS 1977), allowing a shift in the threshold of reporter activity needed to confer the His^+^ phenotype and beneficial for reporters that allowed leaky *HIS3* expression in WT cells (**Fig. 1b**). We reasoned that using two different promoters for each class of TSS-shift reporter would allow us to prioritize mutants more likely to affect promoter scanning globally as opposed to reporter-specific alleles. We previously developed and characterized the *imd2Δ::HIS3* reporter where *HIS3* replaces the *IMD2* ORF and is under control of the *IMD2* promoter (MALIK *et al*. 2017). Downstream shifts to TSS usage bypass a terminator element in the *IMD2* promoter and allow expression of the downstream ORF (KOPCEWICZ *et al*. 2007; JENKS *et al*. 2008; KUEHNER AND BROW 2008). We found this reporter exquisitely sensitive to defects in Pol II and GTFs leading to downstream TSS shifts (MALIK *et al*. 2017) and also that we could identify alleles of *SSL2* (ZHAO *et al*. 2021) or *TFA1*, *TFB3*, or *RPB7* (BASNET *et al*. 2026) able to suppress this reporter through genetic screens or site-directed mutagenesis. Therefore, this reporter appears sensitive to mutations in multiple PIC components that might each alter promoter scanning in distinct ways, *i.e.* controlling Pol II initiation efficiency or controlling scanning processivity.

**Figure 1.**
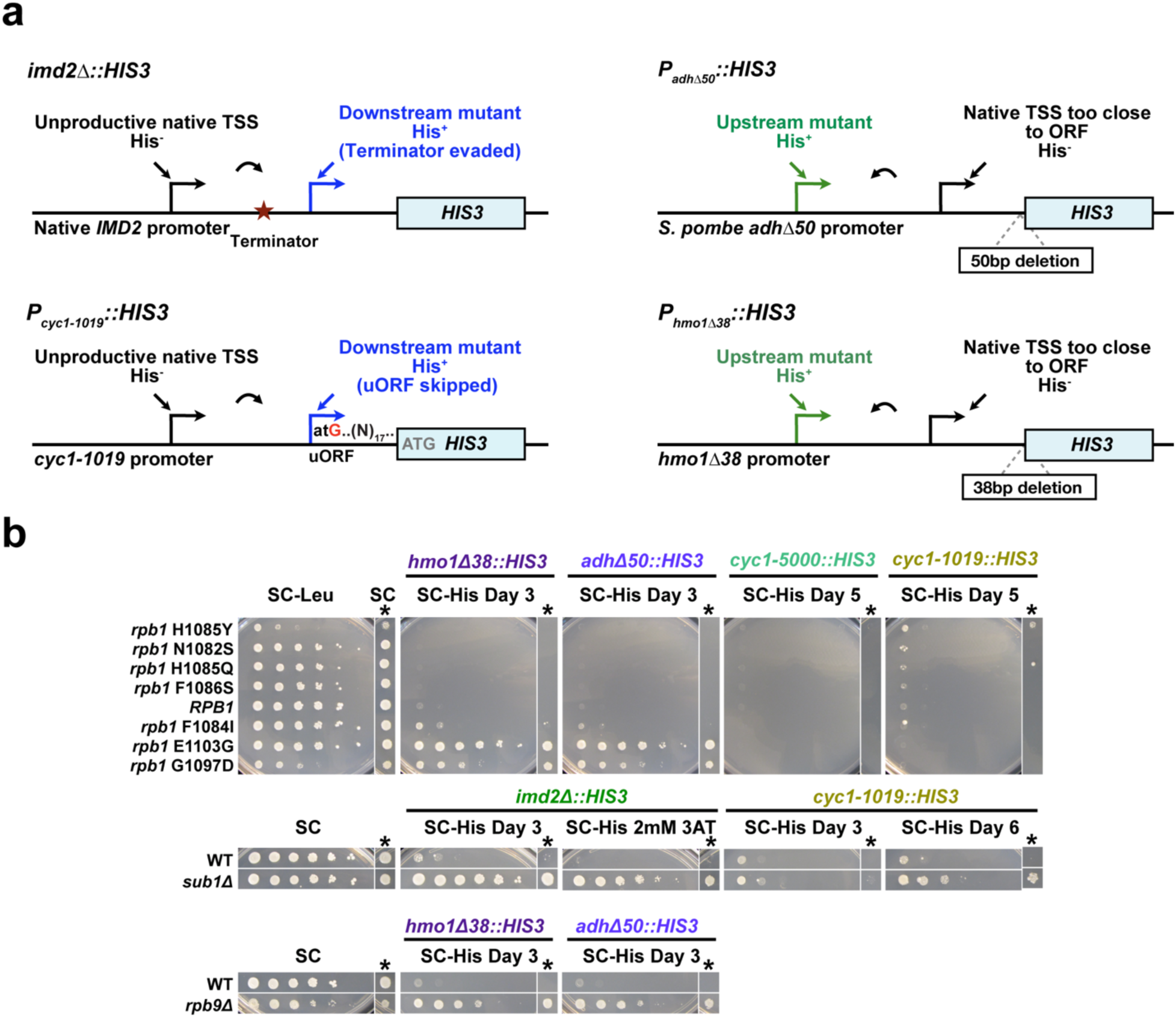
TSS shift-sensitive reporters detect TSS shifts by *HIS3* expression. a. Left. Schematic of initiation reporters *imd2Δ::HIS3* and *cyc1-1019::HIS3* sensitive to downstream shifts in TSS usage. Downstream-shifting TSS mutants constitutively express *IMD2* transcripts from a downstream TSS that allows functional *HIS3* expression when *HIS3* replaces the *IMD2* ORF, conferring a His^+^ phenotype (top). Downstream TSS shifts at *cyc1-1019::HIS3* allow an upstream AUG to be bypassed, resulting in usage of the appropriate in-frame AUG, allowing functional His3p translation (bottom). **Right.** Schematic of the initiation reporters *adhΔ50::HIS3* (top) and *hmo1Δ38::HIS3* (bottom) that are sensitive to upstream shifts in TSS usage. Truncation places the native TSSs for these promoters too close to the native AUG rendering ineffective translation initiation. Upstream shifts in TSS selection lengthen the 5′ UTRs for each reporter, allowing His3p to be translated from the lengthened *HIS3* mRNA. **b.** Validation of reporter phenotypes with known TSS-shifting mutants: 10-fold serially diluted yeast strains or a single concentration of cells (marked with asterisk – “single spot phenotyping”) are plated on media to illustrate reporter function. Here, the *cyc1-5000* allele-based reporter was deployed for isolating *sua* alleles, where the *cyc1-5000* allele resembles *cyc1-1019* allele (1a, left), possessing additional C→A and C→G mutations at -23 and -17 positions respectively, relative to +1 A of the native ATG. This makes the context flanking the upstream AUG start codon more preferred, making the *cyc1-5000* downstream shift detection more stringent than the *cyc1-1019* reporter. *rpb1* alleles (upstream or downstream shifts), *sub1Δ* (downstream shift), and *rpb9Δ* (upstream shift) mutants were tested against the upstream and downstream shift sensitive reporters to illustrate allele-specific reporter activity. Note that *cyc1-1019::HIS3* can be expressed by strong downstream shifting *sub1Δ* and *rpb1* H1085Y of all downstream *rpb1* mutations tested.

Our other downstream shift-detecting reporter *cyc1-1019::HIS3* functions analogously to *imd2Δ::HIS3* (NA *et al*. 1992; PINTO *et al*. 1992a; PINTO *et al*. 1992b). Here, downstream shifts in TSS usage allow bypass of an introduced upstream ATG (the *cyc1-1019* allele). The introduced ATG allows translation initiation out of frame with *HIS3* and confers a His^-^ phenotype. Shift to use of downstream TSSs allows translation to begin at the normal *HIS3* ATG. A variant of this reporter configuration (*cyc1-5000*) was used to identify the first *SUA7* (TFIIB) alleles (PINTO *et al*. 1992a; PINTO *et al*. 1992b) as well as alleles of *RPB1* (*SUA8*) (BERROTERAN *et al*. 1994). This reporter is less leaky than *imd2Δ::HIS3* and thus more tightly His^-^ even in the absence of 3-AT (**Fig. 1b**). We demonstrated that strong downstream shifting mutants *rpb1* H1085Y and *sub1Δ* were His^+^ using this reporter (**Fig. 1b**).

Our upstream shifting reporters function analogously to each other. In each case, a reduced TSS to ATG distance likely hampers translation initiation of *HIS3* because the *HIS3* ATG is too close to the 5′ end of the mRNA (FURTER-GRAVES AND HALL 1990). 5′ extension of the mRNA through usage of an upstream TSS allows more efficient *HIS3* translation initiation. We employed *adh50Δ::HIS3*, previously shown to detect *rpb9* alleles as upstream shifters (FURTER-GRAVES AND HALL 1990; FURTER-GRAVES *et al*. 1991), and a second, novel upstream reporter. To create an additional upstream shift-detecting reporter, we analyzed our previous genome wide TSS data (QIU *et al*. 2020) for promoters that showed especially large shifts in TSS usage in response to upstream TSS mutants, settling on *HMO1*. We then fused different lengths of the *HMO1* promoter, truncated by 38, 45 and 53 basepairs to *HIS3* to ask if any of these configurations could cleanly detect upstream TSS-shifting mutants (**Fig. S1a**). First, we tested if these variants could function effectively as potential initiation reporters by assessing if they showed suppressibility when exposed to UV mutagenesis (**Fig. S1a**), an approach previously tested for reporter suppressibility (HAMPSEY 1991). After observing reporter suppressibility, we tested them in conjunction with TSS-shifting Pol II and *rpb9* mutants (**Fig. 1b**).

### Identification of novel upstream and downstream TSS-shifting mutants

For our selections, we integrated our engineered reporters (*cyc1-1019::HIS3*, *adhΔ50::HIS3* and *hmo1Δ38::HIS3*) at a gene-free locus on chromosome I (**Fig. S1b**) (GRUBER *et al*. 2012) with *imd2Δ::HIS3* present at the native *IMD2* locus. We isolated suppressors of initiation reporters by genetic selection of His^+^ candidates and did so in two batches with two versions of genetic selection strains. In an initial pilot genetic selection, ∼700 candidates were isolated across the four first generation strains carrying the reporters described above, with the majority isolated from the *IMD2* reporter. Candidates were re-phenotyped for their His^+^ phenotypes and for secondary conditional or transcription-related phenotypes. These included growth on YPD at 30 °C, 16 °C, 37 °C and with 3% formamide, SC supplemented with MPA to detect MPA sensitivity commonly observed for known upstream shifting TSS alleles (ZHAO *et al*. 2021), SC-Lys medium to test for suppression of the Spt^-^ reporter allele *lys2-128∂* (SIMCHEN *et al*. 1984; CUI *et al*. 2016), and YPRafGal with YPRaf as a control for suppression of *gal10Δ56* (GREGER AND PROUDFOOT 1998; GREGER *et al*. 2000; KAPLAN *et al*. 2005), a reporter allele able to detect different kinds of transcription-associated phenotypes. We utilized profiles of phenotypes to potentially cluster mutants by similarity of their additional phenotypes and to choose candidates from distinct phenotypic groups for further study. Additionally, known phenotypes of existing mutants allowed provisional attribution of candidates to possible candidate genes. We chose 168 of these candidates for sequencing using a custom Illumina AmpliSeq amplicon pool (WANG *et al*. 2025). Of 168 candidates sequenced using Illumina AmpliSeq, we identified a potential causative mutation in 72 of them. For genes already linked to TSS selection, we identified alleles of *SUA7* (2 unique alleles), *RPB1* (8 unique alleles), *RPB2* (8 alleles from 11 candidates), *RPB9* (2 unique alleles), and *SUB1* (3 alleles from 4 candidates). Because of inconsistent coverage with AmpliSeq, we piloted whole genome sequencing for 10 of the initial candidates where AmpliSeq failed to identify a candidate mutation. Whole genome sequencing (WGS) identified potential causative mutations for 7/10 candidates sequenced and provided information allowing refinement of our selection procedures as described below. This preliminary analysis allowed us to refine our selections for a second round where we identified additional both upstream- and downstream-shifting candidates (**Figs. 2, 3, 4, 5**). We identified transcription-related mutants (subset displayed in **Figs. 2, 3, 4, S2, S3**) but also identified classes of mutant that appear reporter specific (*e.g.* guanine metabolism, **Fig. 5a**) and a likely indirect modulator of TSS selection (e.g. *PMR1*, **Fig. 5b**), classes that AmpliSeq would have failed to identify. Candidates mutations identified by sequencing are compiled in **Table S1.** Previously identified TSS shifting alleles are compiled in **Table S3**. We then incorporated secondary screening to filter these latter classes out as discussed in the next section. Among upstream-shifting candidates, we filtered out recessive loss of function mutants in *RPB9* by a complementation assay. In a subsequent third round genetic selections we isolated ∼1750 more candidates comprising 447 suppressors of *imd2Δ::HIS3*, 672 of *cyc1-1019::HIS3* (**Fig. 2a**), 293 of *adhΔ50::HIS3,* and 350 of *hmo1Δ38::HIS3* (**Fig. 3a**) and after phenotyping-based filtering we prioritized ∼250 candidates for whole genome sequencing. In a separate selection of *imd2Δ::HIS3* suppressors performed by the students in the BIOSC 0068 Foundations of Biology Laboratory 2 course at the University of Pittsburgh, 48 His^+^ candidates were prioritized by phenotypic and genotypic screens for whole genome sequencing, yielding 46 candidates. These were combined with the above 447 *imd2Δ::HIS3* suppressors amounting to 495 *imd2Δ::HIS3* suppressors (**Fig. 2a**). Prioritization across all reporter classes resulted in a total of ∼300 candidates sequenced by whole genome sequencing. Out of the ∼300 candidates sequenced, we identified reporter transcription-associated mutations in ∼240 of these and further ∼160 unique mutants.

**Figure 2.**
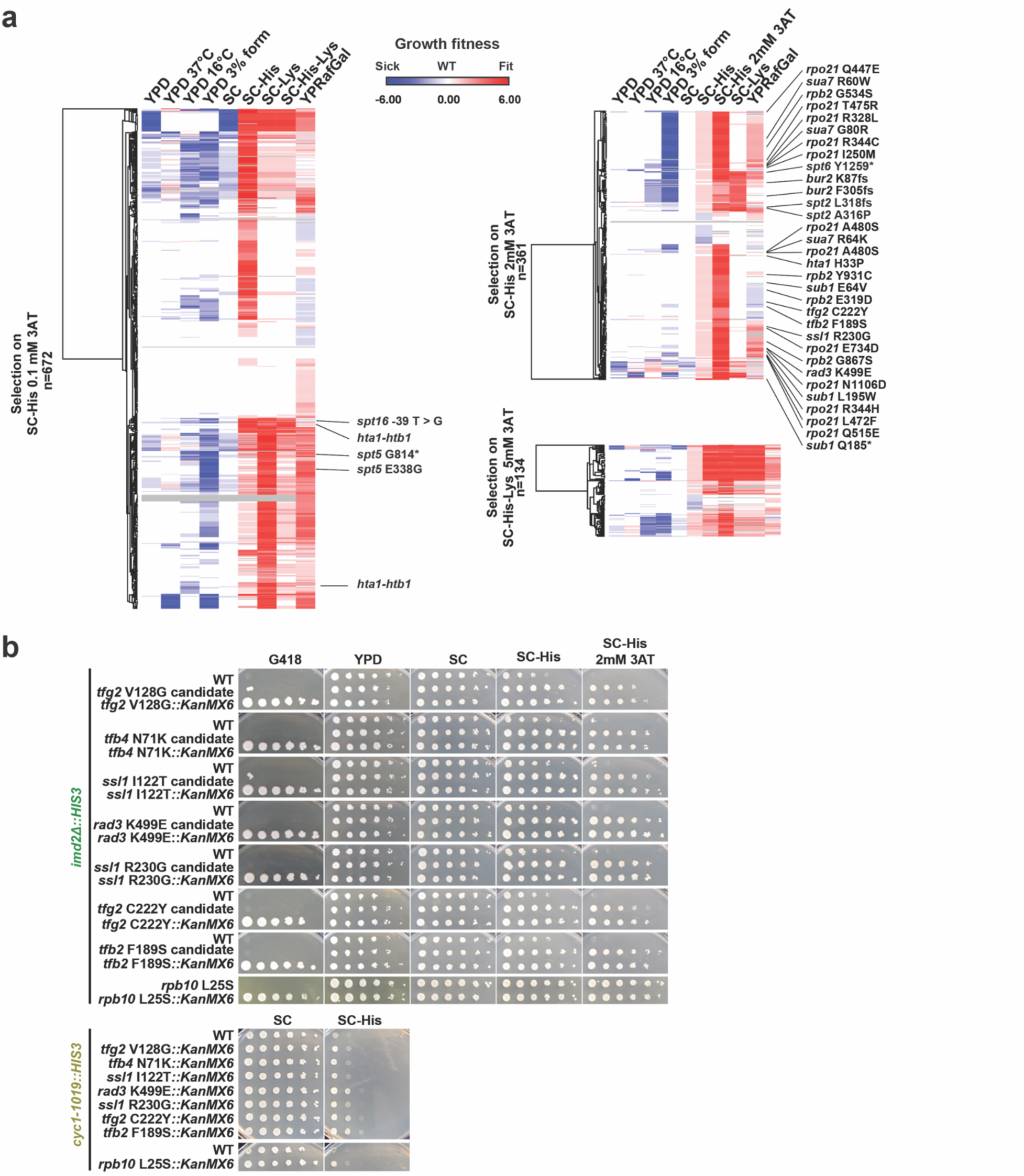
Isolation of spontaneous suppressors of downstream TSS-shift sensitive initiation reporters. **a**. Hierarchically clustered growth phenotypes for a subset of candidate downstream-shifting suppressors of *cyc1-1019::HIS3* (left) or *imd2Δ::HIS3* (right) reporters with a subset of identified mutations annotated; right bottom panel displays *imd2Δ::HIS3* suppressors isolated by selection on SC-His-Lys+5mM 3AT. Phenotypes of candidate suppressors displayed are relative phenotypes of candidates that have been normalized to control conditions, as assessed from single-spot phenotyping assays. Phenotypes were scored from spot assay, corrected to WT and then corrected to the same on control condition (see Methods). Phenotypic quantification therefore assigns a value of zero for WT-like growth, a negative value if phenotype is worse than WT and positive, if growth is better than WT and value of phenotype represents strength of defect. **b.** Validation of a subset of candidates by knock-in of TSS mutants into *imd2Δ::HIS3* and *cyc1-1019::HIS3* reporter backgrounds and re-phenotyping. Spot assay phenotyping of candidates isolated from genetic selection (candidate) and reincorporated mutants with a *kanMX6* tag (*corresponding mutation::kanMX6*) displaying replication of candidate phenotypes in reincorporated mutants. Mutations were amplified out of isolated candidates, PCR sewed with a *kanMX6* drug marker cassette and re-introduced into each of the reporter strains by integrative transformation. The *rpb10* candidate mutation was introduced using CRISPR/Cas9.

**Figure 3.**
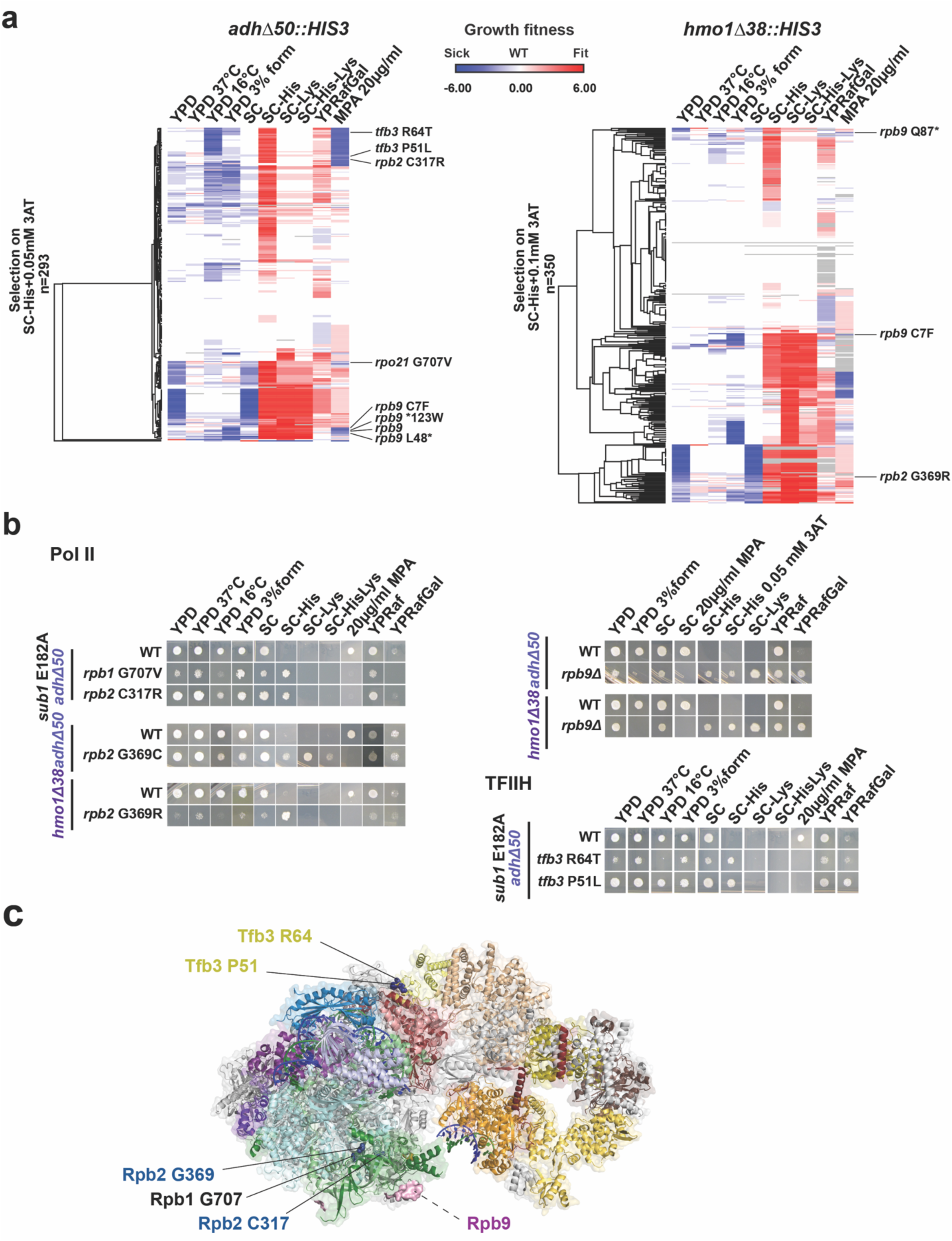
Isolation of spontaneous suppressors of upstream TSS-shift sensitive initiation reporters. **a.** Hierarchically clustered single-spot phenotypes of a subset of upstream-shifting suppressors of *adhΔ50::HIS3* (left) and *hmo1Δ38::HIS3* (right) initiation reporters with identified mutations annotated. Phenotypes displayed have been scored and processed as in Fig. 2a. Briefly, scored phenotypes have been corrected to WT on the same condition, followed by normalization to the same on control conditions. Here, a positive value represents growth better than WT and a negative value represents worse than WT (zero). **b.** Single spot phenotyping of upstream isolated mutants. **c.** Cryo-EM structure of Pol II PIC annotated with upstream shifting mutation residues identified (PDB: 7O4J).

**Figure 4.**
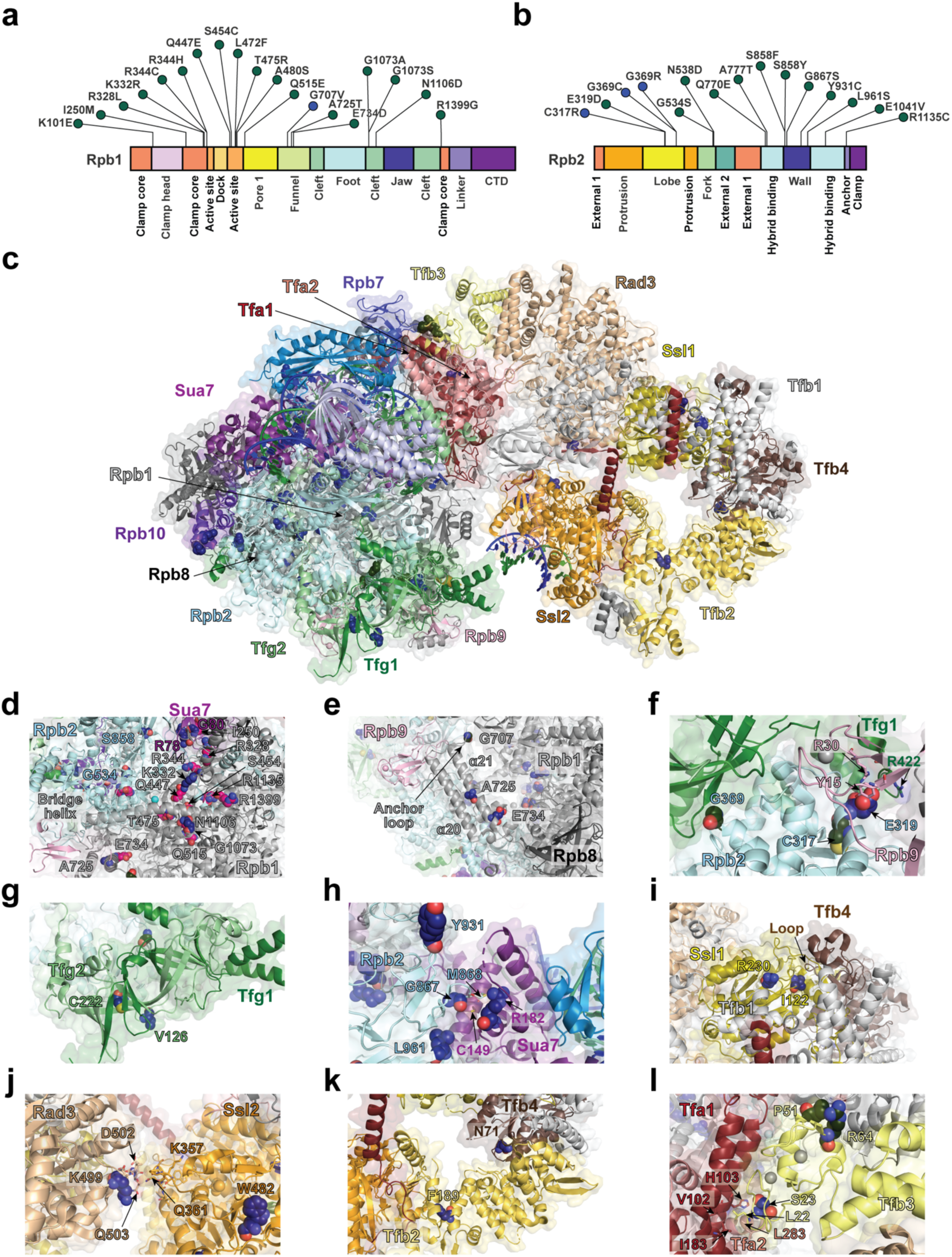
Mutations identified across PIC. Lollipop diagram of mutations isolated in Rpb1 (**a**) and Rpb2 (**b**). Domain diagram figure created using IBS 2.0 (XIE *et al*. 2022). **c**. PIC structure (PDB: 7O4J) displaying mutations isolated in our study. **d.** Structural mutations surrounding active cleft that are formamide sensitive marked in magenta. **e.** Mutations in ⍺21 helix and anchor loop isolated with proximity to Rpb9 and Rpb8. **f.** Residues at Rpb2-TFIIF-Rpb9 interface found to alter TSS selection. **g.** Initiation mutations near Tfg1-Tfg2 interface causing downstream shifts. **h.** Mutations altering interactions between Sua7 B-core and Rpb2 isolated in this study in the vicinity of previously isolated Sua7 C149 initiation mutant. **i.** Potential interactions between Ssl1 and Tfb4 loop or Tfb1 in driving TSS selection. **j.** Rad3 K499E mutation potentially driving TSS selection by altering interactions with Ssl2 across Rad3-Ssl2 interface with K357R and K357E previously isolated in our study altering TSS selection. **k.** Mutations in Tfb2 F189 and Tfb4 N71 lying within the respective structural domains potentially promoting structural “breathing”. **l.** Mutation S23F in the RING domain of Tfb3 increasing interaction between TFIIH and TFIIE by potentially promoting allele-specific hydrophobic packing.

**Figure 5.**
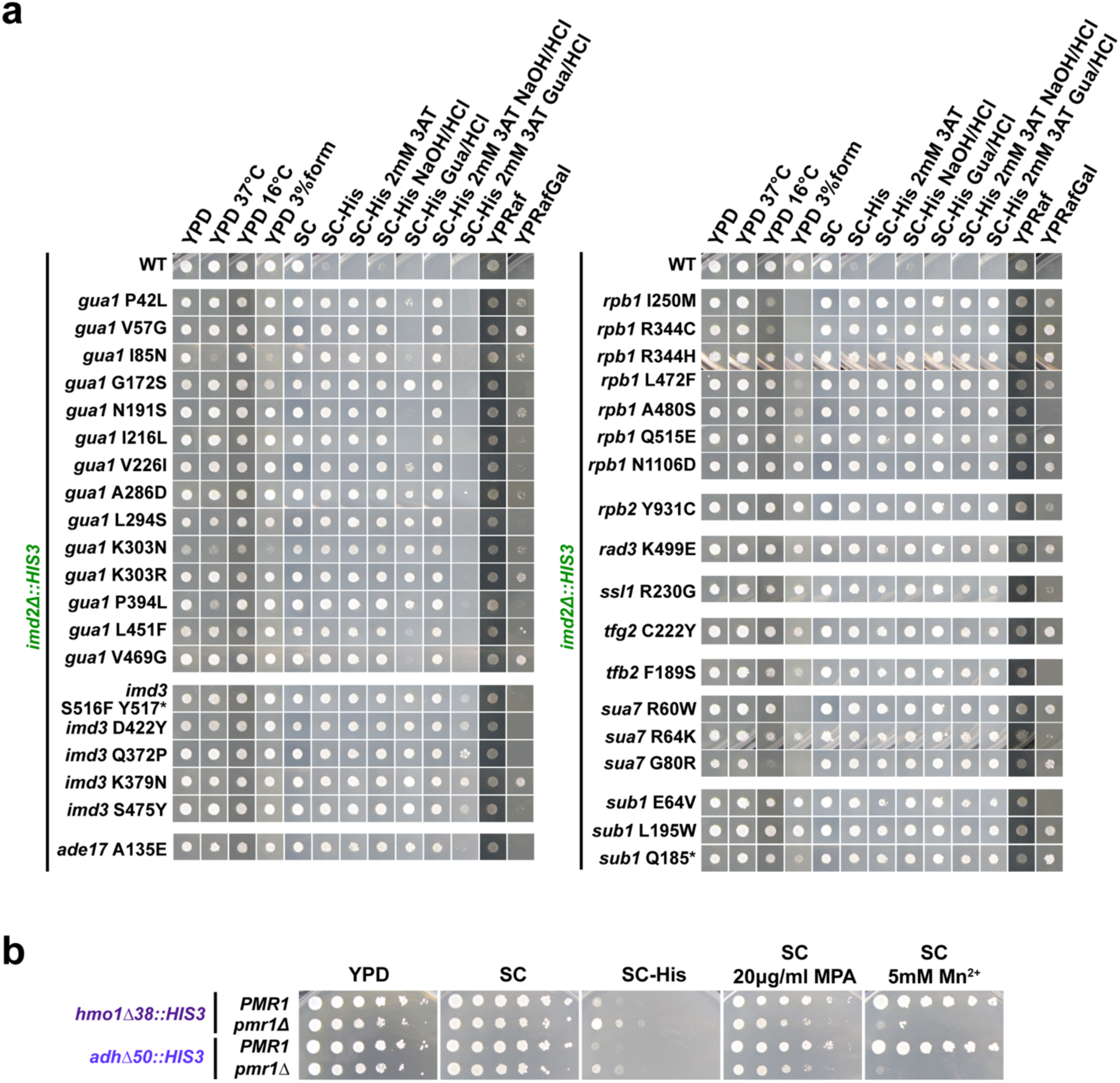
Phenotyping of subset of non-PIC mutants identified as suppressors of initiation reporters. **a.** Guanine metabolism mutants and controls plated as single concentrations of cells on phenotyping media. Guanine metabolism gene mutants confer the His^+^ phenotype to cells with the *imd2Δ::HIS3* reporter, likely due to the *IMD2* promoter’s sensitivity to guanine levels. Guanine supplementation suppresses the His^+^ phenotype of guanine pathway mutants (SC-His 2 mM 3AT Gua/HCl: left panel) whereas PIC mutants remain His^+^ under the same conditions (SC-His 2 mM 3AT Gua/HCl: right panel). Guanine is solubilized in 10 mM NaOH at a final concentration of 500 µM and media must be neutralized by equivalent HCl as NaOH alone affects solubility of MPA. **b.** 10-fold serial dilutions of *PMR1* or *pmr1Δ* strains plated on phenotyping media. *pmr1Δ* suppresses both upstream reporters and confers sensitivity to Mn^2+^.

Of all transcription-related mutants isolated in this study, we observed three primary groups: PIC mutants, indirect effectors of TSS selection (*e.g. PMR1*) or promoter specific non-transcription factors *e.g.* guanine mutants), and chromatin/Pol II elongation factors. We validated 8 candidates through reconstruction of mutants by generating alleles in a clean genetic background and all 8 recapitulated candidate suppression (**Fig. 2b**). In addition to direct validation by reconstruction of a subset of mutants, several lines of evidence support the mutations identified by WGS are causative for phenotypes. First, we selected spontaneous suppressors, so the expectation was for most strains to have essentially a single causative mutation. Second, we identified previously described alleles of *SUA7*, *RPB1*, *RPB2*, *TFB3*, *SPT6*, and *TFA1* while not observing changes elsewhere in their genomes. Third, we identified multiple inactivating alleles of *SUB1* and *RPB9* as expected for non-essential genes with known effects on TSS selection, while not observing changes elsewhere in their genomes. Fourth, there were five cases where we identified the identical allele independently in separate selections. Finally, nearly all candidate sequencing data showed only a single change in a gene that could rationally be linked to transcription initiation. Because previous genetic interaction studies have helped annotate functions to mutations (JIN AND KAPLAN 2014; ZHAO *et al*. 2021), we tested inter-subunit interface mutants and genetic interactions across a subset of our mutants before assessing TSS shifts caused by our novel mutants genome wide, which we discuss later.

Among mutants in PIC components, we identified alleles of Pol II (*RPO21/RPB1*, *RPB2*, *RPB9*, *RPB10*) and of GTFs (TFIIB, *TFA1*/TFIIE, *TFG2*/TFIIF, TFIIH: *SSL2*, *SSL1*, *TFB2*, *TFB3*, *TFB4* and *RAD3*) (**Fig. 4**). We first observed a high prevalence of mutations in the downstream shifting class in *RPB1* and *RPB2* lining the cleft leading to and surrounding the active site. Because *RPB1* and *RPB2* are larger genes, these mutants predominated in our selections, though many of these mutations were novel (**Figs. 4a, b**). Because downstream shifting is expected from reduced Pol II activity (KAPLAN *et al*. 2012; MALIK *et al*. 2017), catalytic or structural defects that lead to reduced catalytic activity or PIC function are expected to be isolated as downstream shifters. However, not all *rpb* mutants clustered around the active site (**Figs. 4a, b**). We observed two types of mutations amongst those surrounding the activity cleft. We found that in both Rpo21/Rpb1 and Rpb2, mutations within structural domains surrounding the cleft displayed formamide sensitivity (**Fig. 4d**), a phenotype that could reflect protein folding or stability defects. Mutations facing or on the surface of the cleft generally were formamide resistant. We also isolated mutations at intra-subunit interfaces including those linked to control Pol II catalysis. For example, we isolated an upstream-shifting mutation G707V in the Rpb1 anchor loop region (**Fig. 4e**). This loop lies between ⍺20 and ⍺21 helices of Rpb1 (KASTER *et al*. 2016) and a poly-alanine substitution at Rpb1 707-709 in the anchor loop was previously shown to confer MPA sensitivity, consistent with an upstream shift in TSS usage (KASTER *et al*. 2016). We additionally observed mutations in the Rpb1 ⍺21 helix, A725T and E734D, both resulting in downstream shifts. A prior mutation in Rpb1 ⍺21, G730D, was identified as a reduced catalytic activity mutant and appeared to function through incompatibility with Rpb9 in that slow growth of G730D was suppressed by *rpb9Δ* (KASTER *et al*. 2016). It is possible that these substitutions work similarly though they are predicted to be more subtle. Alternatively, E734 is positioned to potentially interact with a network of residues that connects to an Rpb1-Rpb8 interaction but is also adjacent to TFIIS when it is inserted into the active site. However, no role for TFIIS in TSS selection has yet been described. Regardless, perturbation to E734 could result in reduced Pol II catalysis, leading to a downstream shift in TSS usage.

Of *rpb2* mutations, beyond the active cleft, we observed mutations in Pol II-lobe-jaw domain along the path of DNA entry near the Rpb9-TFIIF interface (HEMMING *et al*. 2000) (**Fig. 4f**). Because previous studies have shown that perturbation of both TFIIF and Rpb9 functions results in upstream shifts (FURTER-GRAVES *et al*. 1991; HULL *et al*. 1995; SUN AND HAMPSEY 1995; SUN *et al*. 1996; KHAPERSKYY *et al*. 2008; EICHNER *et al*. 2010), the upstream-sensitive reporter suppressor Rpb2 C317R could be consistent with disruption of this interface. Here, Rpb2 C317 is buried, and substitution of an arginine might be predicted to perturb structure and alter positioning of either Rpb9 or TFIIF or both. Disruption of Tfg1 at this interface encompassing residue R422 (homologous to R417 in the studied *S. mikate* Tfg1) (EICHNER *et al*. 2010) and deletion of *RPB9* (WOYCHIK *et al*. 1991) each result in a cold sensitive phenotype, a phenotype shared by Rpb2 C317R (**Fig. 3b**). In contrast to Rpb2 C317R, a nearby substitution E319D results in the opposite phenotype – suppression of a downstream-shift sensitive reporter. While Rpb2 E319 is close to Tfg1, it appears to make direct interactions with Rpb9 Y15 and R30 and perturbing the length of the 319 sidechain in E319D may alter this interaction with Rpb9 (**Fig. 4e**).

We also isolated Rpb2 G369C and G369R mutants as upstream shifters (**Fig. 4f**). Substitutions in Rpb2 G369 were previously isolated as Spt^-^ (HEKMATPANAH AND YOUNG 1991), as a suppressor of a TFIIB downstream-shifting allele (CHEN AND HAMPSEY 2004), and to be MPA^S^ with altered termination at a genetic reporter allele involving *ADH2* (KUBICEK *et al*. 2013). Bulky substitutions for glycine at this residue would be in position to interfere with the Rpb1-Tfg1 interface (CHEN *et al*. 2007; SCHILBACH *et al*. 2021). Previous studies show that mutation in the TFIIF dimerization domains within both Tfg1 and Tfg2 subunits also resulted in upstream TSS shifts (SUN AND HAMPSEY 1995; GHAZY *et al*. 2004; EICHNER *et al*. 2010). Intriguingly, we also isolated a downstream shifting mutation in the Pol II subunit Rpb10, a subunit shared amongst the three eukaryotic nuclear RNA polymerases. This allele is discussed below where we probe how this substitution may be working (**Fig. 6**).

**Figure 6.**
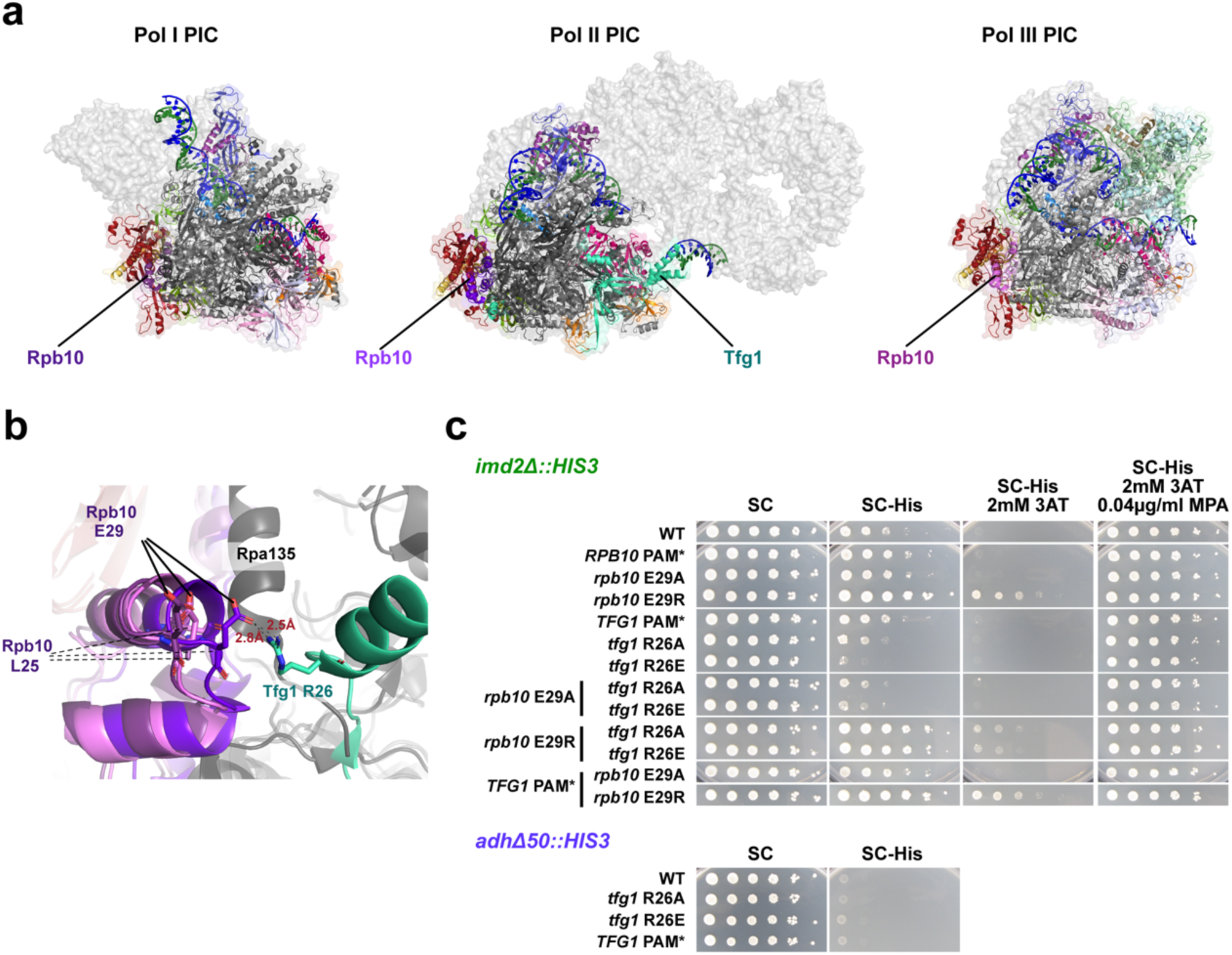
Analysis of potential Rpb10-Tfg1 interactions in proper TSS selection. **a.** Cryo-EM structures of PICs of Pol I, Pol II, and Pol II displaying Rpb10, a shared subunit between them. **b.** Cryo-EM structure of Rpb10 from Pol I-III PICs (Pol I PIC: 6TPS, Pol II PIC: 7O4J, Pol III PIC: 6EU0) overlayed displaying Tfg1 from the Pol II PIC, with Tfg1 R26 in the proximity of Rpb10 E29 suggesting a potential electrostatic interaction. While Rpa135 in Pol I lies adjacent to Rpb10, a specific physical interaction with Rpb10 E29 may not exist. **c.** Serially diluted spot assay phenotyping of single and double mutants of *tfg1* R26 and *rpb10* E29 resides displaying charge reversals, mutations to alanine and silent PAM control mutations (Rpb10 L30 AAC→AAT) as mutants were generated by CRISPR/Cas9.

Among GTFs, we identified novel alleles of *TFG2* (TFIIF), *SUA7* (TFIIB)*, RAD3, SSL1, SSL2, TFB2*, *TFB4* and *TFB3* (TFIIH) and *TFA1* (TFIIE). For Tfg2, we identified three suppressors of a downstream-shift sensitive reporter: V128G, C222Y (**Figs. 2b**, **4g**) and T366P (**Table S1**) (with C222Y isolated twice). This is the first occurrence of *tfg2* alleles potentially conferring downstream shifts, because previously identified mutants of Tfg2 that affected TSSs resulted in upstream shifts (SUN AND HAMPSEY 1995; GHAZY *et al*. 2004; EICHNER *et al*. 2010). Tfg2 V128 and C222 are close to each other in space but there is not an obvious interpretation for their structural perturbations. Interestingly, while Tfg2 substitutions V128G and C222Y lie within the Tfg2 dimerization domain (but not at the dimerization interface), T366P lies in the winged helix domain (WH) and is not visualized in the yeast PIC. The WH domain has been proposed to interact with upstream DNA (EICHNER *et al*. 2010).

In *SUA7,* we isolated novel suppressors of downstream-shift sensitive reporters in the B-finger/B-reader domain such as R60W and G80R alleles as well as those in previously studied residues R64K (BANGUR *et al*. 1997; PARDEE *et al*. 1998) and R78C (PINTO *et al*. 1994) (**Fig. 4h, Table S1**). We note that G80R is cold sensitive just as are classical *sua7* alleles (PINTO *et al*. 1992b). Besides substitutions in the B-reader domain, we also identified R182C in the Sua7/TFIIB B-core N-terminal cyclin fold (**Fig. S2**), the domain that interacts with the Pol II wall, involved in directing the DNA-RNA hybrid (CRAMER *et al*. 2001; KOSTREWA *et al*. 2009). The TFIIB B-core domain binds to the Pol II complex at the Rpb2 “wall” domain, which undergoes reorganization as a consequence of the TFIIB-Pol II interaction. Previously isolated substitutions in C149 at the TFIIB/Rpb2 interface resulted in a downstream TSS shift (WU *et al*. 1999). Our new substitution, Sua7 R182C is in a residue that packs with Sua7 C149, which is adjacent to the Rpb2 wall domain residue M868 (KOSTREWA *et al*. 2009; SAINSBURY *et al*. 2013) (**Fig. 4h**). We also isolated Rpb2 substitutions in the wall near the N-terminal B-cyclin fold domain proximal to M868 (Rpb2 G867S, Y931C, L961S). Unbiased isolation of TSS mutants at this interface highlights the ability of our selections to identify interfaces involved in TSS selection.

Within TFIIH, we isolated alleles across the complex, consistent with a network of interactions throughout its subunits able to modulate the scanning process. Within Ssl1, *ssl1* I122T and *ssl1* R230G were isolated and are the first *ssl1* alleles directly connected to TSS selection. *SSL1* was initially genetically isolated through a suppressor of a stem loop placed at the 5′ UTR of *HIS4* (YOON *et al*. 1992) and potentially regulates Rad3 helicase activity involved in NER (COIN *et al*. 1998; KIM *et al*. 2015) via its VWA family protein domain. Rad3 activity is not implicated in TFIIH initiation functions and the location of our *ssl1* alleles suggests that they might instead alter internal Ssl1 interactions or the organization of residues that interact with Tfb1 or Tfb4 or both. Ssl1 I122 is directly adjacent to a loop of Tfb4 that inserts into Ssl1 while Ssl1 R230 is positioned internally to organize an Ssl1 loop that connects to both Tfb1 and Tfb4 interfaces (**Fig. 4i**).

Further, we isolated Tfb2 F189S that lies in the HEAT repeats domain of Tfb2 as a suppressor of a downstream shift-sensitive reporter. Tfb2 F189 contributes to hydrophobic packing within Tfb2 that might limit dynamics or breathing of this critical connector protein for bridging Ssl2 to the rest of TFIIH. Relaxation of this domain through loss of the bulky phenylalanine sidechain potentially could confer flexibility on the connection of Ssl2 to the rest of the complex (**Fig. 4k**). Similarly to how Tfb2 connects Ssl2 to the rest of TFIIH, Tfb4 is the connector of Tfb2 to the rest of TFIIH. We isolated *tfb4* N71K as a suppressor of a downstream shift-sensitive reporter. Tfb4 N71 lies in the VWA domain and insertion of a bulkier and charged lysine into an internal Tfb4 pocket might also loosen the Tfb4 structure, potentially adding flexibility to the linkages across components (**Fig. 4k**).Tfb3 is critical connecting TFIIH by its RING domain to Pol II (through Rpb7) and the rest of the PIC (through TFIIE). Previously tested mutations that disrupt this interface cause upstream shifts in TSS selection (YANG *et al*. 2022; YANG *et al*. 2024; BASNET *et al*. 2026), while removal of the Tfb3 RING domain or the TFIIK kinase module (part of TFIIH) result in usage of TSSs ∼30 bp downstream of the TATA box *in vitro*, similarly to non-scanning eukaryotes (MURAKAMI *et al*. 2015; YANG *et al*. 2022). We have interpreted these mutants as conferring a defect in scanning, either a loss of scanning processivity or reduction in scanning rate. Here, we have isolated *tfb3* S23F as suppressing a downstream shift-sensitive reporter (**Fig. 4l**). Tfb3 residue S23 lies within the Tfb3 RING finger domain and is present in a loop adjacent to a hydrophobic interface with TFIIE subunits Tfa1 and Tfa2, formed by Tfb3 L22, Tfa1 I183, Tfa1 V102, Tfa1 H103, and Tfa2 L283 (**Fig. 4l**). Intriguingly, a potentially more disruptive substitution at the very same residue, *tfb3* S23P, was isolated as an upstream shifter along with other disruptive substitutions at the Tfb3-TFIIE interface (YANG *et al*. 2022; YANG *et al*. 2024; BASNET *et al*. 2026). Because it has the opposite phenotype of Tfb3 S23P, we interpret Tfb3 S23F as stabilizing the Tfb3-TFIIE interaction (see Discussion). Supporting our interpretations of mutations identified in targeted *TFB3* and *TFA1* genetic screens or site-directed mutational studies (YANG *et al*. 2022; YANG *et al*. 2024; BASNET *et al*. 2026), we isolated mutations R64T and P51L in *TFB*3 that result in suppression of upstream shift-sensitive reporters, namely Tfb3 R64T and P51L (**Fig. 3, 4l**). Tfb3 R64 had previously been identified as making a key interaction with Rpb7 and each side of this interaction appears to function in promoting scanning as mutations are MPA^S^ and/or alter TSS selection (BRABERG *et al*. 2013; YANG *et al*. 2022; BASNET *et al*. 2026). Contemporaneously with our identification of these residues in unbiased genetic selections, P51 mutations were directly tested based on evolutionary analysis of *TFB3*, with P51 and R64 conferring strong MPA-sensitivity. Because P51 and R64 substitutions were epistatic with each other, this result was consistent with these residues working together, *e.g.* P51 working by enabling positioning of R64 (YANG *et al*. 2024).

For TFIIH, physical interactions between Rad3 and Ssl2 may also contribute to TFIIH function in scanning (**Fig. 4j**). Rad3 K499E resulted in suppression of a downstream-shift sensitive reporter and is directly proximal to Ssl2 (**Figs. 2b**, **4j**). We had previously identified mutations in Ssl2 at K357R and K357E (ZHAO *et al*. 2021) near this Rad3 interface sharing the same phenotype as Rad3 K499E, raising the possibility that could be functional interface, with the caveat that Ssl2 K357 may act through other interactions. Because we interpret downstream shifts for Ssl2 mutants as a gain of processivity, there are two possibilities for Rad3 K499E in this framework. First, the interaction between Rad3 and Ssl2 might limit Ssl2 processivity, and therefore we would interpret K499E as a loss of function for the interaction. Alternatively, the interaction between Rad3 and Ssl2 might promote processivity, and K499E confers phenotypes by strengthening the interaction.

We further wanted to isolate special “*ssl2*-like” alleles that showed both a His^+^ phenotype on the downstream shift-sensitive *imd2Δ::HIS3* reporter and a Lys^+^ phenotype due to suppression of the *lys2-128∂* Spt^-^ phenotype reporter, a phenotype combination exhibited by a specific subset of downstream-shifting *ssl2* mutants (ZHAO *et al*. 2021). These previously identified alleles were highly localized to the DRD domain of Ssl2 (ZHAO *et al*. 2021). Before our Ssl2 work, the Spt^-^ phenotype had only been observed in conjunction with upstream-shifting phenotypes for alleles of Pol II and the TFIIF subunit Tfg2 (KAPLAN *et al*. 2012; JIN AND KAPLAN 2014). Because the Spt^-^ *ssl2* mutants appeared novel, we wanted to ask if additional GTF mutants with similar phenotypes could be identified, therefore we performed selections with the *imd2Δ::HIS3* reporter selecting on SC-His-Lys medium. While we identified two new alleles of *SSL2*, each with a mutation in Helicase domain 1 (HD1, *ssl2* R378L and *ssl2* W482G), we did not identify additional mutations of this class in other GTFs, suggesting this class of allele may be unique to *SSL2*. Finally, we isolated a TFIIE mutation *tfa1* E374* as a putative downstream shifting allele (**Fig. S2**). Tfa1 has been shown to have inhibitory effect on Tfa2-activated Ssl2 translocase activity (LIN AND GRALLA 2005). This suggests *tfa1* E374* could function by relieving restriction on Ssl2 translocase activity, resulting in downstream TSS shift due to increased processivity.

### Non-PIC mutants conferring suppression of TSS-shift sensitive reporters

Besides PIC factors, our study also isolated non-PIC mutants, comprising putative reporter-specific factors like guanine pathway mutants, indirect effectors like *PMR1*, and chromatin/transcription elongation factors. We found almost fifty percent of initial *IMD2* reporter suppressors had mutations in guanine metabolism genes (**Fig. 5a**). *IMD2* is activated under low GTP conditions, specifically because low GTP induces a TSS switch that bypasses the upstream terminator in the *IMD2* promoter (**Fig. 1a**). Therefore, defects in guanine metabolism create inducing conditions for *IMD2.* We predicted this class of mutant might be differentially sensitive to supplementation by guanine. We found that guanine supplementation suppressed the His^+^ phenotype of guanine pathway mutants specifically, but not a subset of tested PIC mutants (**Fig. 5a**), and therefore incorporated guanine supplementation into subsequent phenotyping conditions as indicated above. Our WGS also identified an allele of *PMR1* as a suppressor of the *adhΔ50::HIS3* reporter (**Table 1S**). *PMR1* encodes an ER/golgi Ca^2+^/Mn^2+^ transporter, and loss of its activity raises cytoplasmic Mn^2+^ levels, conferring Mn^2+^ hypersensitivity (RUDOLPH *et al*. 1989; DURR *et al*. 1998; MANDAL *et al*. 2000) (**Fig. 5b**). Our lab had previously found that *pmr1Δ* genetically interacts with increased activity Pol II mutants (BRABERG *et al*. 2013), that increased activity Pol II mutants are specifically sensitive to increased Mn^2+^ (CABART *et al*. 2014), and that *pmr1Δ* itself resulted in an upstream shift to TSS usage at the *ADH1* promoter (QIU *et al*. 2016), due to potential increased Pol II activity in response to heightened cytoplasmic (and nuclear) Mn^2+^ levels. Isolating *pmr1* alleles as suppressors of the *hmo1Δ38::HIS3* reporter would be consistent with potential global effects of Mn^2+^ levels on TSS selection. While Mn^2+^ has many effects on Pol II transcription including perturbing fidelity, it consistently stimulates RNA polymerase catalysis *in vitro e.g.* (NIYOGI *et al*. 1981; QIU *et al*. 2024). Therefore, upstream shifts at multiple promoters *in vivo* are consistent with expectations for increased Pol II catalytic activity during initiation caused by increase in cellular Mn^2+^ levels.

Intriguingly, we isolated chromatin-related Pol II associated elongation factors as suppressors of both downstream reporters *imd2Δ::HIS3* and *cyc1-1019::HIS3* (**Fig. S3**). Many of these are canonical “Spt” genes and fittingly, some of these were isolated in our double selections for His^+^ Lys^+^ (Spt^-^) suppressors, but a number were isolated from His^+^ selections. We isolated alleles of *spt2*, *spt5*, *spt16*, *bur2, spn1* and histones from both downstream-sensitive reporters and *spt4* and *spt6* only from the *imd2Δ::HIS3* reporter (**Fig. S3**). This reporter bias may potentially exist because our selections were not saturating. A role for Spt6 in *IMD2* TSS selection has been described for specific *spt6* alleles defective for interaction with a phosphorylated linker in Rpo21/Rpb1 (CONNELL *et al*. 2022). The chromatin/elongation factors we isolated are all known to suppress initiation from cryptic promoters within genes (WINSTON *et al*. 1984; CLARK-ADAMS *et al*. 1988; KAPLAN *et al*. 2003; CHEUNG *et al*. 2008; DORIS *et al*. 2018). Similarly, their Spt^-^ phenotypes at *lys2-128∂* are consistent with a chromatinized-promoter within a transcription unit (in this case, a Ty1 ∂ element within the *LYS2* gene) being activated upon elongation-mediated chromatin disruption in specific mutants (SIMCHEN *et al*. 1984; WINSTON *et al*. 1984; CLARK-ADAMS *et al*. 1988; SWANSON *et al*. 1991; PRELICH AND WINSTON 1993; FISCHBECK *et al*. 2002; KAPLAN *et al*. 2003; CHEUNG *et al*. 2008; CUI *et al*. 2016; DORIS *et al*. 2018). This would be a reasonable model to explain suppression of *imd2Δ::HIS3* because the *IMD2* promoter (driving His^+^) is transcribed as part of an upstream CUT transcript (KOPCEWICZ *et al*. 2007; JENKS *et al*. 2008; KUEHNER AND BROW 2008), potentially sensitizing the downstream TSS to chromatin defects driven by Pol II elongation. Conversely, this model does not easily explain why we also identified chromatin/elongation gene alleles as suppressors of *cyc1-1019::HIS3* (see Discussion) and raises the possibility that chromatin factors might globally alter TSS selection, which we test directly below.

### Functional characterization of TSS mutants through genetic interaction analyses

After our genetic selections, we wanted to functionally characterize mutants using genetic interactions. In our prior studies, both additive/synergistic and epistatic interactions have been observed for TSS-shifting mutants leading to a framework where mutants are interpreted as affecting initiation efficiency (i.e. Pol II catalysis) or scanning processivity (TFIIH/PIC functions). Previous examination of double mutants between Pol II catalytic mutants and TFIIB or TFIIF alleles showed additivity across combinations (enhancement when mutants shift TSSs the same direction and suppression when they shift in opposite directions). However, in other combinations, we observed epistasis where double mutants exhibited the TSS phenotype of one of the single mutants and not the other. This was especially clear when upstream shifting Pol II mutants were combined with downstream shifting *ssl2* alleles, leading to the hypothesis that the tested *ssl2* alleles increased scanning processivity, but this didn’t lead to downstream TSS shifts in mutants that increased initiation upstream. Regardless of mechanistic models for how mutants are genetically interacting, double mutant analysis can reveal when mutations are dependent on one another (epistasis), which is indicative of their defects being at the same step, or when mutations function redundantly in an essential step (synergy or synthetic lethality), or when mutations are perturbing a system independently, whereby double mutant phenotypes will be the additive consequences of the single mutants combined. To therefore functionally dissect our mutants, we tested interactions at inter-subunit interfaces and further analyzed functional interactions across the PIC (**Figs. 6b, 6c; 7; S4**).

We first tested a potential Rpb10-Tfg1 interface in TSS selection by genetic interaction studies and site-directed mutagenesis. We isolated *rpb10* L25S as a downstream-shifting suppressor of the *imd2Δ::HIS3* reporter and showed this allele could also suppress the *cyc1-1019::HIS3* reporter (**Fig. 2b**). Because Rpb10 is a common subunit shared amongst three RNA polymerases, it was interesting to find a Pol II-specific phenotype. Examination of Pol PIC structures for Pol I, II, and III PICs suggested that an interaction between Rpb10 in the vicinity of the Rpb10 helix-loop-helix L25 residue might only be present in the Pol II PIC, where Rpb10 appears to contact a loop in the Tfg1 N-terminal domain (**Fig. 6b**). This potential interaction suggested that Rpb10 might affect Pol II TSS distributions through an interaction with TFIIF subunit Tfg1 that may be perturbed in the *rpb10* L25S allele. To examine this, we targeted Rpb10 E29 and Tfg2 R26 residues that display potential interactions based on structural studies (SCHILBACH *et al*. 2021) (**Fig. 6c**). We generated a series of genomic mutations by CRISPR including charge reversals at both residues and assessed growth and TSS-shift reporter phenotypes. On analysis of mutants, we did not observe overall growth defects in *tfg*1 single or *tfg1 rpb10* double mutants (**Fig. 6c**, rows 6 and 7). Charge reversal *rpb10* mutation of E29R resulted in suppression of the downstream shift reporter while E29A did not display effects (**Fig. 6c**, rows 3 and 4). However, when *rpb10* E29R was combined with the *tfg1* R26A or R26E mutations, apparent suppression of the downstream shift phenotype was observed (**Fig. 6c**, rows 10 and 11). Moreover, we found that *tfg1* mutants also suppressed the leaky His^+^ phenotype of *imd2Δ::HIS3*, with Tfg1 R26E being more robust than R26A (**Fig. 6c**, rows 6 and 7, SC-His panel). Suppression of the leaky His^+^ phenotype of *imd2Δ::HIS3* might result from an upstream TSS-shifting phenotype, therefore we tested these *tfg1* mutants against the upstream reporter *adhΔ50::HIS3* (**Fig. 6c**, lower panel) but *tfg1* mutants did not suppress *adhΔ50::HIS3*. These results show that an additional *rpb10* allele can alter TSS selection (**Fig. 6c**) and that its phenotype can be modulated by proximal *tfg1* alleles. However, we could not prove that Rpb10-Tfg1 interactions underlie the *rpb10* L25S mutant phenotype.

We tested additional functional interactions by tetrad dissection analysis of diploids followed by fitness analysis of each spore by pixel quantification using EBImage (PAU *et al*. 2010; NAMJILSUREN AND ARNDT 2025). We then calculated “expected” double mutant growth phenotype from single mutant phenotypes by a multiplicative model for independent effects on fitness of single mutants in a double mutant combination (MANI *et al*. 2008) and compared this expected double mutant growth phenotype to the observed double mutant growth phenotype. Genetic interactions were established based on comparisons of these double mutant metrics. We examined upstream shifting *rpb9*Δ genetic interactions with a panel of mutants. First, we examined *rpb9*Δ in conjunction with the upstream-shifting *rpb2* G369C as the proximity of Rpb9 to Rpb2 and TFIIF raises the possibility that these mutants work in the same pathway, as it has previously been proposed that *rpb2 G369C* works through effects on TFIIF. Second, we tested *rpb9Δ* against a subset of our novel downstream shifting alleles to ask if they were additive or epistatic. The GTF TFIIF binds at the interface between Rpb9 and Rpb2 in the protrusion-lobe domain with proximity to Rpb2 G369 (**Fig. 4, 7b**)(CHEN *et al*. 2007). Because *rpb2* G369C and *rpb9Δ* are both upstream shifting mutations, it is expected that the double mutant in case of non-interaction would be a product of individual fitnesses. Conversely, if they functioned together in the same process, we might observe less than multiplicative effects (sometimes referred to as a specific form of epistasis). Instead, we observed growth fitness of *rpb2* G369C *rpb9Δ* double mutants to be worse than predicted from the multiplicative model, indicating a synthetic defect (**Fig. 7d**).

**Figure 7.**
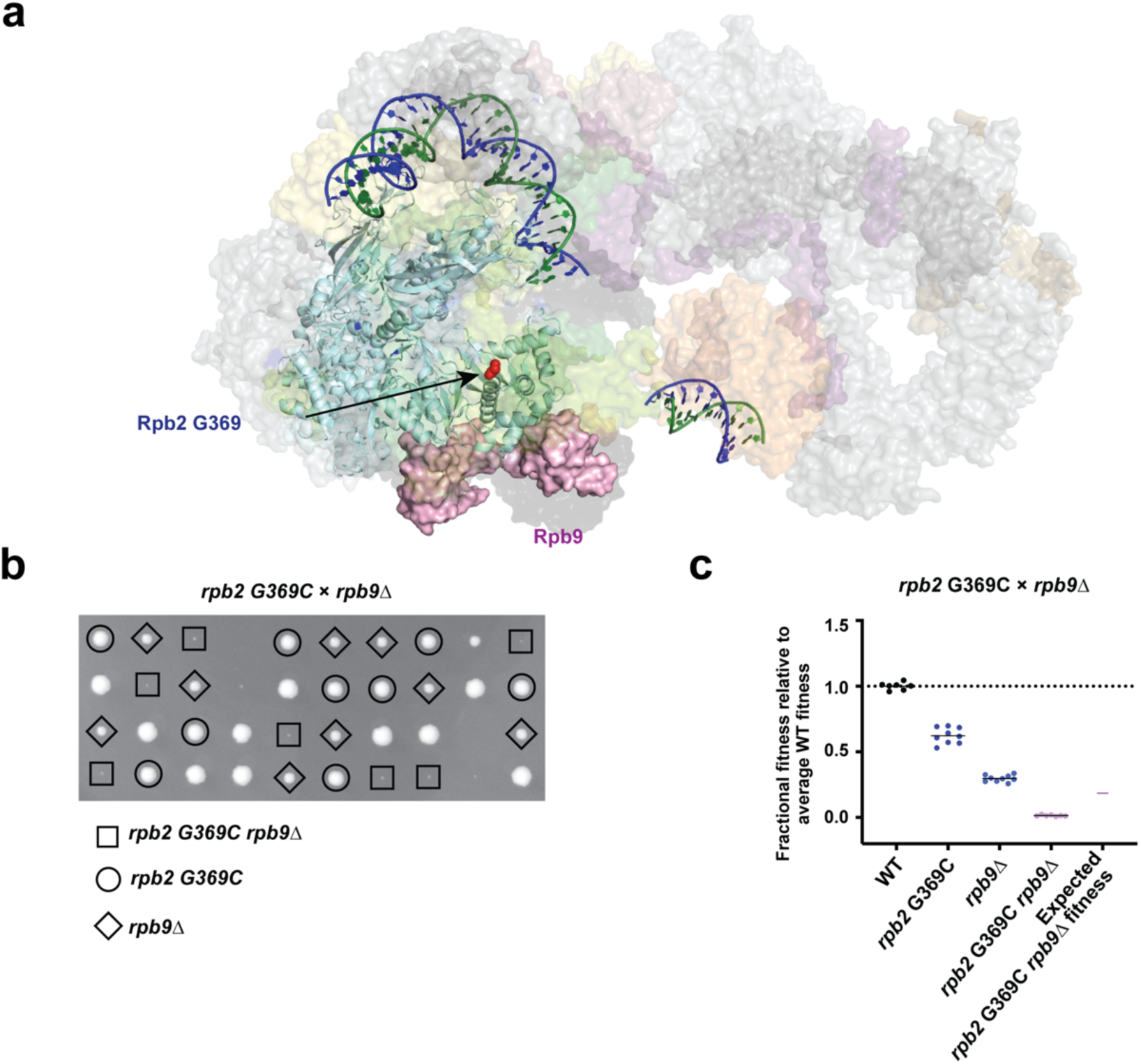
Synthetic genetic interaction between *rpb9Δ* and *rpb2* G369C. **a.** Cryo-EM structure of Pol II PIC displaying Rpb2 G369 and Rpb9. Rpb2 G369 lies in the Pol II lobe domain which interacts with Rpb9 though this region between Rpb2 and Rpb9 and has been shown to interact with TFIIF. **b.** Tetrad dissection of *rpb9Δ/+ rpb2* G369C/+ diploid to examine genetic interactions between them. Spores are annotated by genotype. **c.** Observed and expected fitness of single and double mutants plotted for assessing genetic interaction. “Expected” fitness for a double mutant is the product of the two single mutant fitnesses (MANI *et al*. 2008). Observed fitness of single and double mutants estimated by quantification of spore colony sizes. Observed fitness is worse than expected fitness indicating a synthetic genetic interaction.

When combining upstream shifting *rpb9Δ* with a panel of downstream shifting mutants, *tfb2* F189S, *tfb4* N71K, *ssl1* I122T, *rad3* K499E, *rpb10* L25S and *tfg2* C222Y by tetrad dissection analysis (**Fig. S4**). We found *rpb9Δ rpb10* L25S double mutant fitness to be similar to the *rpb9Δ* single mutant growth defect, with resultant fitness around the predicted fitness (**Fig. S4**, bottom, middle). In addition, we found *rpb9Δ tfg2* C222Y double mutant fitness to be slightly worse than expected (**Fig. S4**, bottom, right). Because these mutants have opposite effects on TSS shifts, combination might be predicted to result in suppression of growth defects, if the effects of each mutant on TSS selection are additive *and* the growth defects observed correspond to the TSS selection phenotype. This kind of interaction between *rpb9Δ* and *rpb10* L25S to be epistatic where the defect of one mutation (upstream) cannot be compensated by a second mutation working in an opposite direction (downstream). Conversely, for interaction between *tfg2* C22Y and *rpb9Δ*, if either of the *tfg2* C222Y or *rpb9Δ* effects required a WT allele for their specific phenotypes, then instead of suppression of phenotypes by two opposite acting alleles, we might observe exacerbation due to conversion of a downstream or upstream shifting allele to the other class in the double mutant. Assuming the change in growth fitness reflects an equivalent change in TSS selection, this kind of interaction could represent sign epistasis. In contrast to the *rpb9Δ-tfg2* C222Y negative interaction, we observed clear double mutant suppression between *rpb9Δ tfb2* F189S, *rpb9Δ tfb4* N71K, *rpb9Δ ssl1* I122T and *rpb9Δ rad3* K499E, where each TFIIH allele suppressed growth defects associated with *rpb9Δ* (**Fig. S4**, top panel and bottom, left). This indicates that the growth defect induced by upstream mutant *rpb9Δ* was compensated by all tested TFIIH downstream mutants and suggesting that the growth defect of *rpb9Δ* could be initiation related. This supposition is because TFIIH’s gene expression functions are limited to the PIC. This result was unexpected based on a simple model where the *rpb9Δ* upstream-shifting phenotype would be caused by loss of its restraint of the key Pol II catalytic domain, the trigger loop (TL) (KASTER *et al*. 2016). This model was based on *rpb9Δ* conferring similar phenotypes to hyperactive mutants in the Rpb1 TL (upstream TSS shifts and increased elongation rate *in vitro*). Upstream-shifting TL mutants are epistatic to downstream-shifting *ssl2* TFIIH alleles, and therefore similar interactions between *rpb9Δ* and downstream-shift reporter specific-TFIIH alleles was what was previously expected (see Discussion).

### Novel PIC alleles shift TSSs genome wide while chromatin/elongation alleles have muted effects

To gain insight into the function of a subset of our TSS shifting mutants, we quantified TSSs across a curated set of promoters genome wide by STRIPE-seq in WT and mutant strains (POLICASTRO *et al*. 2020; POLICASTRO *et al*. 2021; POLICASTRO AND ZENTNER 2022) (**Fig. 8a**). We chose 8 downstream-shifting mutants, six from the PIC class and two from the chromatin/elongation class isolated in our studies namely *tfb4* N71K, *rad3* K499E, *ssl1* I122T, *tfb2* F189S, *rpb1* A480S, and *rpb1*0 L25S (PIC mutants) and *spn1* V199G and *spt5* G814* (chromatin/elongation mutants) together with their corresponding WT parents (**Figs. 8**, **9**, **S5**, **S6**). To assess TSS shifts in mutants relative to WT parents, we quantified TSSs as in our previous studies (QIU *et al*. 2020; ZHAO *et al*. 2021) along with STRIPE-seq processing as in (POLICASTRO *et al*. 2021). We focused our TSS analyses to 5979 Pol II promoter regions (RHEE AND PUGH 2012; QIU *et al*. 2020; ZHAO *et al*. 2021) filtered to include promoters passing a cutoff of a read count threshold in the promoter window analyzed resulting in analysis of n=2829 promoters. TSS usage values were then compared to those in WT by two TSS shift metrics as in (QIU *et al*. 2020; ZHAO *et al*. 2021): shift in median TSS (Δmedian), which represents the difference in median position of the TSS distribution at a promoter between a mutant and WT strains (**Fig. 8b**) and change in spread (Δspread) that represents the change in width of promoter TSS distributions relative to WT (**Fig. 9a**).

**Figure 8.**
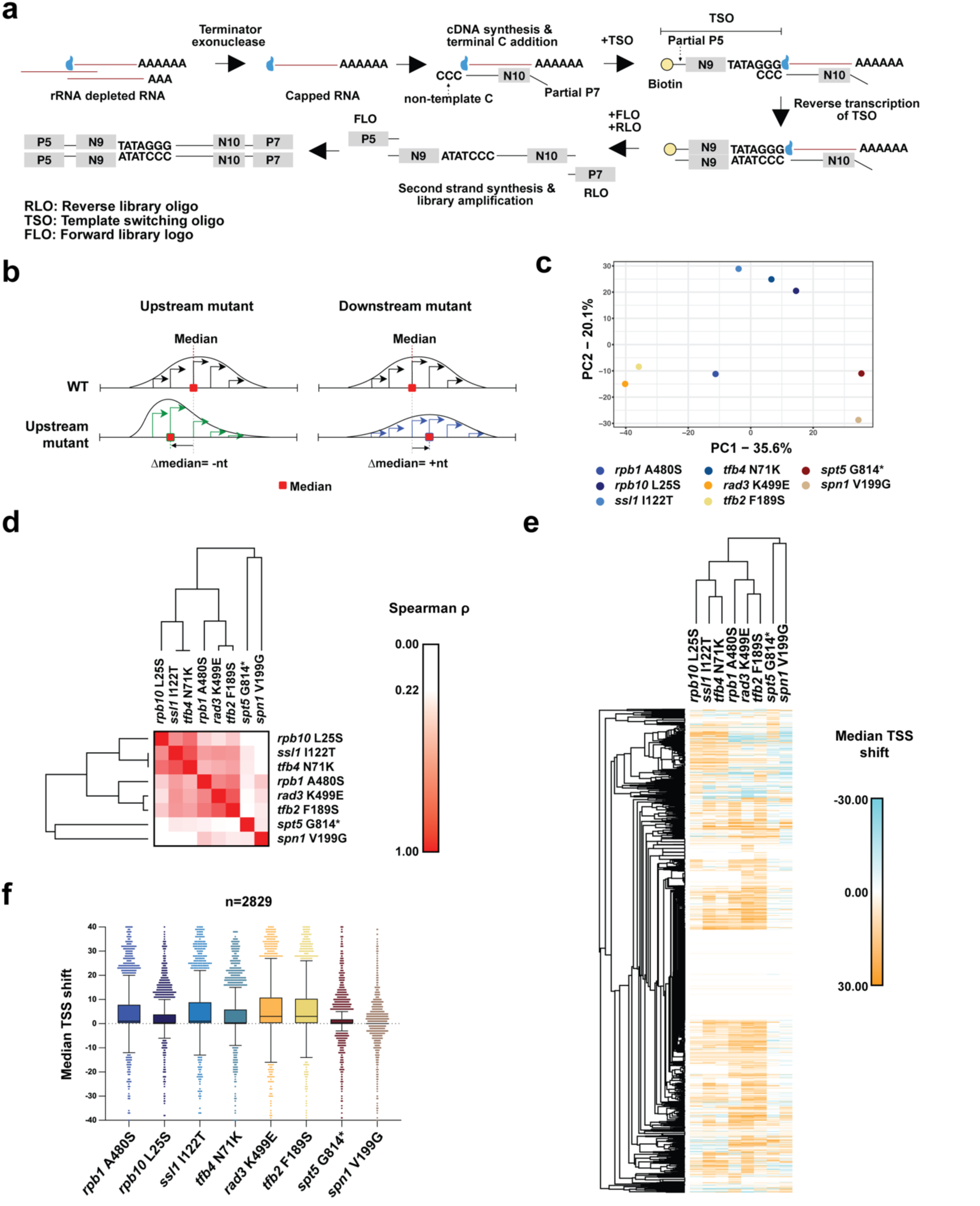
STRIPE-seq analysis of novel mutants identifies three functional groups of initiation mutants based on shift of median of TSS distribution. **a.** Schematic of STRIPE-seq library generation. **b.** Schematic of Δmedian metric for TSS analysis. **c.** Principal component analysis (PCA) of Δmedian values of the top 2829 promoters based on an expression cutoff of ≥10 reads within the analyzed 401 nucleotide promoter windows. PCA of Δmedian values across mutants with combined biological replicates plotted along the first two principal components display three clusters, ① chromatin mutants *spt5* G814*, *spn1* V199G, ② *rpb1* A480S, *rpb10* L25S, *ssl1* I122T, *tfb4* N71K and ③ *rad3* K499E and *tfb2* F189S. d. Hierarchical clustering of Spearman correlation of Δmedian values across mutants. **e.** Hierarchically clustered heatmap of Δmedian values across top 2829 promoters of mutants with replicates combined. **f.** Box-plot analysis of Δmedian values across mutants presented by Tukey plot.

**Figure 9.**
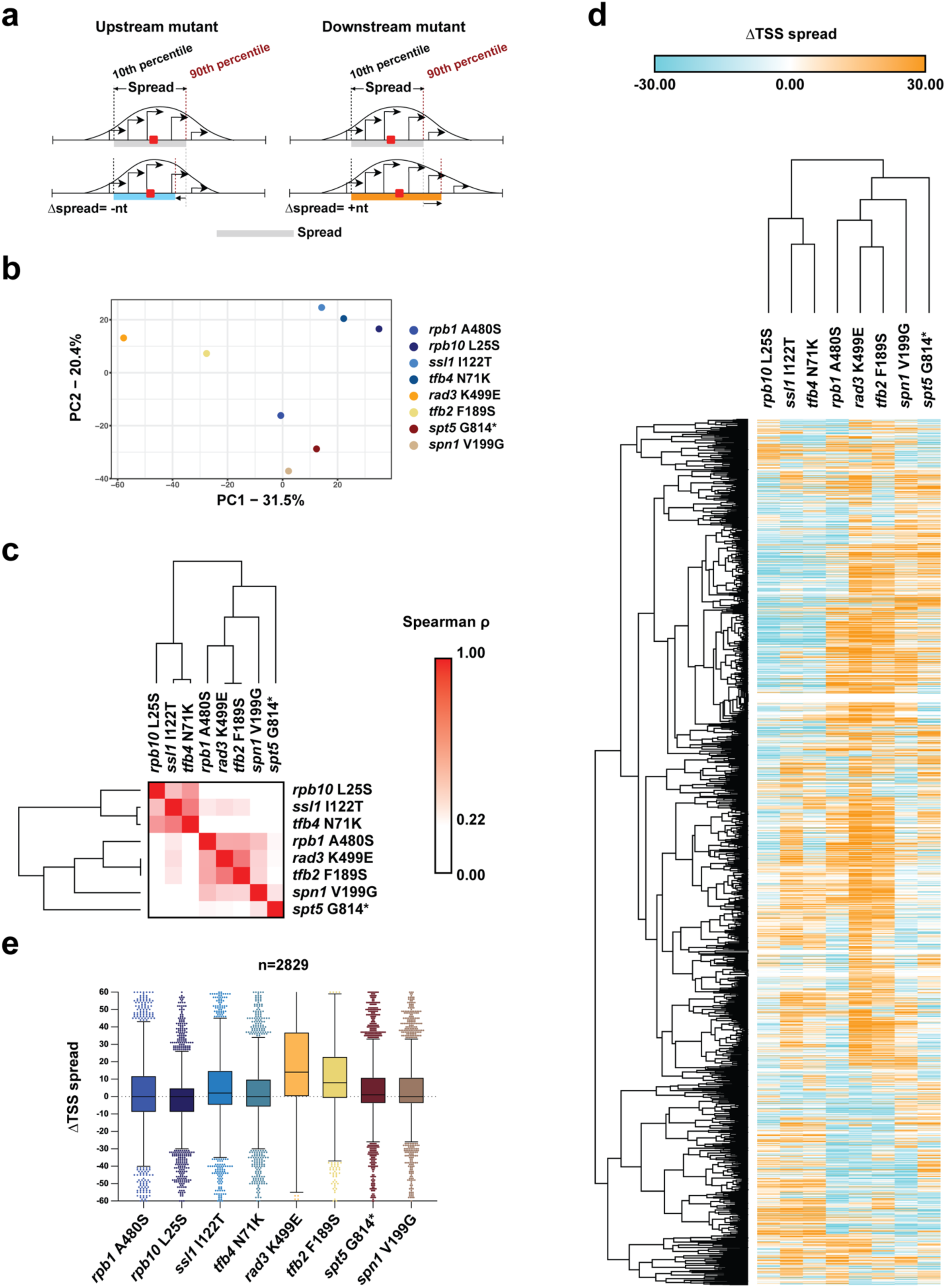
STRIPE-seq analysis of novel mutants identifies three functional groups of initiation mutants based on shift of spread of TSS distribution. **a.** Schematic of Δspread metric of analysis **b.** Principal component analysis of Δspread values of top 2829 genes across mutants plotted along the first two principal components displays three clusters for Δmedian, ① chromatin mutants *spt5* G814*, *spn1* V199G, ② *rpb1* A480S, *rpb10* L25S, *ssl1* I122T, *tfb4* N71K and ③ *rad3* K499E and *tfb2* F189S. c. Hierarchical clustering of Spearman correlation of Δspread values across mutants. **d.** Hierarchically clustered heatmap of Δspread values across top 2829 promoters of mutants. **e.** Box-plot analysis of Δspread values across mutants presented by Tukey plot.

Individual replicates were generally reproducible for Δmedian TSS (**Fig. S5**). Principal component analysis (PCA) of the Δmedian values of combined replicates showed three clusters representing functional differences across the primary axis of variance (**Fig. 8c**): cluster 1 contained *rpb1* A480S, *rpb10* L25S, *ssl1* I122T and *tfb4* N71K, cluster 2 contained *rad3* K499E and *tfb2* F189S, and cluster 3 contained *spt5* G814* and *spn1* V199G. When examining PCA of individual replicates each compared to the average WT median position (**Fig. S5b**), we found chromatin alleles to cluster near the WT, suggesting they only have mild effects on TSS distributions globally, but with greater effects for *spt5* G814* than for *spn1* V199G (**Fig. 8f**). While both chromatin/elongation alleles did have effects beyond just an individual reporter, their effects were quantitatively and qualitatively different from PIC alleles, consistent with potentially relating to the chromatin context of individual promoters rather than innately affecting TSS selection and promoter scanning at all promoters. All PIC alleles showed a large bias towards downstream shifts in TSS distribution across the majority of promoters, with *rad3* K499E and *tfb2* F189S showing the strongest effects (**Fig. 8f**).

PCA of Δspread values also displayed three groups and examination of PCA for individual replicates of all strains compared to the average WT behavior shows that chromatin/elongation mutants cluster with WT, most PIC mutants cluster together while *rad3* K499E and *tfb2* F189S, which showed the strongest effects on Δmedian TSS, formed a separate cluster based on Δspread (**Figs. 9, S6**). Examination of directional bias in spread suggests that while spread was perturbed for most mutants, *rad3* K499E and *tfb2* F189S showed the strongest increase in spread, consistent with the downstream shift in TSS usage also expanding TSS spread to downstream positions (**Figs. 9e, S6d**). These results again differentiated two TFIIH alleles, *rad3* K499E and *tfb2* F189S, from two others, *ssl1* I122T and *tfb4* N71K.

## DISCUSSION

Our studies reveal the wider landscape of genes and specific mutations that can alter the TSS selection in yeast. In addition to independently identifying some already discovered alleles, validating our novel reporters, we also identified an array of novel alleles suppressing our initiation reporters. Notably, we did not find additional factors associated with promoters but not directly connected to the PIC, such as chromatin remodelers or transcription coactivators, as able to suppress our initiation reporters. This could simply be due to these remodelers not strongly functioning in TSS accessibility at our reporters, or at least not in a way that might allow reporter suppression upon their mutation. We find that mutations in many PIC components including Rpb10 and additional TFIIH subunits can perturb initiation by promoter scanning when mutated. Importantly, we find that the novel *rpb10* allele and tested TFIIH alleles alter TSS selection genome wide. Our prior studies, new alleles identified here, and our parallel studies on the Pol II/PIC interface reinforce that upstream shifting of TSSs results from compromise of TFIIH-PIC interactions through the Tfb3-TFIIE-Rpb7 interface (BRABERG *et al*. 2013; YANG *et al*. 2022; YANG *et al*. 2024; BASNET *et al*. 2026). We extend evidence that the Rpb2-TFIIB interaction is important for promoter scanning while also finding that perturbation to Rpo21/Rpb1 and Rpb2 broadly leads to suppression of downstream-shift sensitive reporters. This latter result further supports the idea that defective Pol II catalysis/initiation activity results in phenotypes consistent with downstream shifts in TSS usage. In contrast, we also have identified novel TFIIH downstream-shifting alleles that are generally not located at inter-subunit interfaces, save one.

The downstream-shifting phenotype of these novel TFIIH alleles would be consistent with a gain of function in TFIIH scanning processivity or rate. One interpretation of our alleles in Rad3, Tfb2, Tfb4, and Ssl1 based on their locations is that they alter plasticity or dynamics within the TFIIH structure. When this plasticity is altered, perturbations may propagate either to the Ssl2 active site or to Ssl2 DNA binding properties, which in turn allow scanning to be extended to downstream positions through increased processivity or increased scanning rate. The idea that TFIIH processivity can be sensitive to small structural perturbations provides a conceivable evolutionary path for TFIIH to gain ability to reach downstream TSSs during the advent of the promoter scanning mechanism.

Potential communication between Ssl2 and the rest of the PIC can take a few paths. First, there is a connection from Ssl2 to Tfb3 and Pol II/TFIIE through the following proteins: Ssl2 to Tfb2 to Tfb4 to Ssl1 to Rad3 to Tfb3. Second, Tfb1 drapes over multiple TFIIH components and connects to Tfa1 from TFIIE. Finally, Rad3 has a small number of residues that may close the gap between it and Ssl2 and contact Ssl2 residues. We have identified alleles along components that form the path from Ssl2 to Tfb3 (Tfb2, Tfb4, Ssl1, Tfb3) and between Rad3 and Ssl2. If the protein connection between Ssl2 to Tfb3 has a certain amount of flexibility or dynamics (using an analogy of an accordion’s bellows or a pantograph that might tolerate or allow extension or contraction, depending on rigidity), then alterations to this flexibility might have consequences for Ssl2 rate or processivity as it translocates downstream on promoter DNA. The locations of putative gain of function Tfb2, Tfb4, and Ssl1 alleles are all consistent with perturbing internal intra-subunit interactions and we would propose they increase flexibility of the “bellows” or “pantograph” formed by these components that connect Ssl2 to Tfb3 and the PIC. This increased flexibility may then allow Ssl2 to translocate longer or faster, resulting in a shift in TSS usage to downstream positions. Future biochemical and biophysical experiments, potentially coupled with molecular dynamics simulations, might provide tests of a model that, first, novel TFIIH alleles do indeed confer increased TFIIH processivity, and second, that these alleles might alter structural properties of TFIIH in similar ways.

Chromatin/elongation factors can be envisioned to function in controlling initiation in two ways. First, native chromatin architecture may reduce transcription initiation from nucleosome-occluded TSSs (most yeast promoters have TSSs located within the boundaries of +1 nucleosomes) (RANDO AND WINSTON 2012). If there are downstream shifts in nucleosomes dependent on elongation factors, then this could feed back to TSS usage and relax nucleosomal inhibition of nucleosome-sensitive TSSs for promoters with appropriately positioned +1 nucleosomes. Second, if a promoter’s TSSs are transcribed themselves from an upstream promoter, then promoter-associated nucleosomes that are transcribed dependent on an upstream TSS could become dependent on factors that restore transcribed nucleosomes in the wake of elongating Pol II. This situation might reflect allele-selective effects on TSS usage for transcribed promoters and less so global effects. This arrangement of an upstream promoter driving transcription of a downstream promoter and thereby sensitizing the chromatin over the downstream promoter is similar to that observed at the *SRG1*-*SER3* locus (MARTENS *et al*. 2004) and also the structure of the *lys2-128∂* allele (SIMCHEN *et al*. 1984). Each of these loci are especially sensitive to chromatin and transcription elongation defects, and the latter of which was used in the original identification of a number of the Spt chromatin/elongation factors – factors that we isolated with our reporters here (SIMCHEN *et al*. 1984; CLARK-ADAMS AND WINSTON 1987; CLARK-ADAMS *et al*. 1988; MALONE *et al*. 1991; SWANSON *et al*. 1991; CUI *et al*. 2016). In these cases, chromatin maintenance suppresses transcription initiation from within the transcribed regions. Altered chromatin architecture in chromatin mutants can result in widespread initiation from cryptic promoters (PRELICH AND WINSTON 1993; KAPLAN *et al*. 2003; CHU *et al*. 2007; CHEUNG *et al*. 2008; DORIS *et al*. 2018) but perhaps also from chromatin-sensitive TSSs adjacent to existing promoters. In either case, altered chromatin can result in increased transcription from “suppressed” sites.

We rationalize the identification of chromatin elongation factors from the *imd2Δ::HIS3* reporter because of its idiosyncratic transcription from upstream TSSs (generating a cryptic unstable transcript (CUT) (DAVIS AND ARES 2006; KOPCEWICZ *et al*. 2007; JENKS *et al*. 2008; KUEHNER AND BROW 2008)) that likely contributes to or controls nucleosome positioning over the downstream, low-guanine inducible TSS. It was already known that certain *spt6* alleles could increase usage of downstream TSSs at *IMD2* in the absence of known inducing conditions (CONNELL *et al*. 2022). Here, we extend identification of chromatin and elongation factors that can act at *IMD2* through suppression of *imd2Δ::HIS3* to *spt2*, *spt4*, *spt5*, *spt16*, *spn1*, *bur2, hta1,* and *htb1* alleles. These results suggest that *imd2Δ::HIS3* may be similarly sensitive to the chromatin/Spt^-^ class of mutant as is the classical *lys2-128∂* allele. However, we also isolated chromatin mutants suppressing *cyc1-1019::HIS3* in addition to *imd2Δ::HIS3.* This *cyc1* allele was inserted into a non-native locus that is devoid of genes that has previously been used to house reporter genes (GRUBER *et al*. 2012). Our initial presumption was that the *CYC1* promoter would not be itself transcribed, but examination of data from the Steinmetz lab (XU *et al*. 2009) suggests that our *CYC1* promoter fragment also contains *SUT215*, a stable unannotated transcript, and that integration of *cyc1-1019::HIS3* at chrI:199270-199271 causes it to be flanked by *SUT004* upstream and *CUT442* downstream. Taken together, these provide support that *cyc1-1019::HIS3* sensitivity to chromatin and elongation factors could be through transcription over its promoter altering promoter nucleosomes, and we speculate chromatin/elongation effects at other promoters may relate especially to those promoters that have overlapping transcription units.

### Limitations of the study

Our genetic interaction analysis indicated some interesting relationships between reporter suppressor alleles, specifically the suppression observed between a subset of downstream-shifting TFIIH alleles and *rpb9Δ*. Our previous genetic interaction studies showed that upstream shifting Pol II catalytic alleles were epistatic to downstream-shifting *ssl2* alleles for both growth phenotypes and for TSS shifts at *ADH1* (ZHAO *et al*. 2021). Our previous understanding for *rpb9Δ* was that it was functioning similarly to upstream-shifting Pol II catalytic alleles (KASTER *et al*. 2016). This assumption, coupled with the assumption that novel downstream-shifting TFIIH alleles should function like *ssl2* downstream-shifting alleles, led to the expectation that *rpb9Δ* would be epistatic to TFIIH alleles for TSS shift phenotypes. Additionally, we had the expectation that *rpb9Δ* growth phenotypes related to its elongation defects and would not necessarily be suppressible by initiation factor mutants. Contrary to these expectations, we find the possibility of complexity in genetic interactions that might indicate different initiation alleles are working in more complex ways than we had previously thought. We imagine these results should be developed and extended in a few ways. For example, double mutant combinations of *rpb9Δ* and existing *ssl2* alleles and of Pol II alleles with novel TFIIH alleles would also allow understanding of potential allele specificity. Additionally, examination of double mutants for phenotypes on transcription reporters would allow to assess if initiation specific phenotypes reflect overall growth phenotypes. Finally, quantitative primer extension analysis of double mutants for *ADH1* TSS selection would also determine if growth phenotypes correlate with specific changes to TSS distributions at promoters, where *ADH1* promoter has extensively served as an excellent target for TSS distribution change assessment.

## Data availability

Sequencing data generated in this study are available in the NCBI BioProject under accession number PRJNA1337182. Strains used in this study, outlined in **Tables S1** and **S2**, are available from the corresponding author upon request.

## Acknowledgements

We acknowledge Dr. David Brow at the University of Wisconsin for guidance on genetic selections. We are especially grateful to the students of BIOSC 0068 Foundations of Biology Laboratory 2 course at the University of Pittsburgh for genetic selection of candidates leading to identification of a subset of alleles described here. We acknowledge Gabe Zentner, Robert Policastro, and Chhabi Govind for advice on STRIPE-seq. We would like to dedicate this paper to the memory of Gabe Zentner.

## Funding

Supported by funding from National Institutes of Health grants R01GM120450 and R35GM144116 to C.D.K. This research was supported in part by the University of Pittsburgh Center for Research Computing and Data, RRID:SCR_022735, through the resources provided. Specifically, this work used the HTC cluster, which is supported by NIH award number S10OD028483.

## Conflict of interest

The authors declare no conflicts of interest.

---

**Figure S1.**
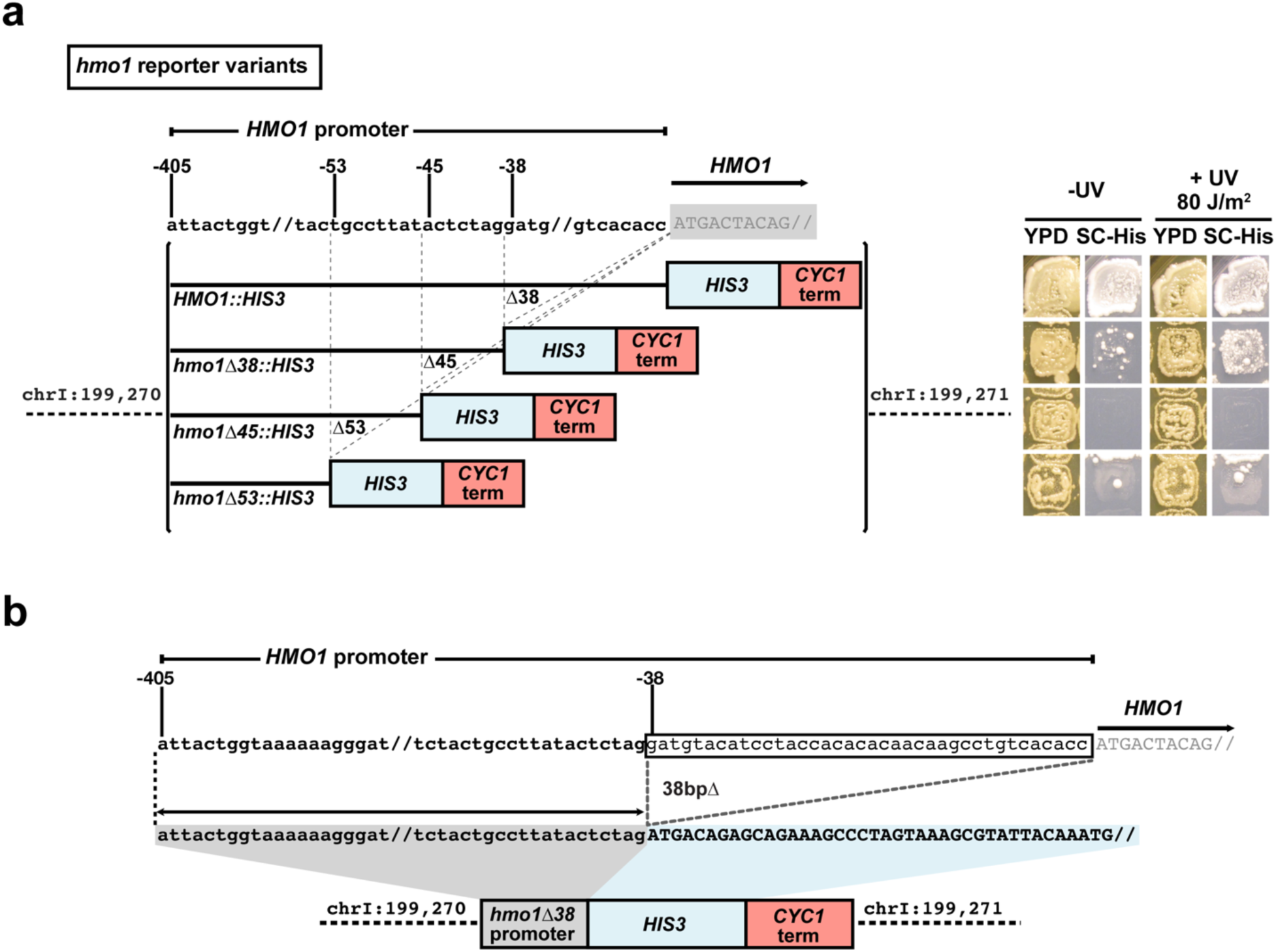
Isolation of *HMO1* promoter variant that detects upstream TSS shifts. **a.** Schematic illustrating an *HMO1* promoter variant deletion series (left) with behavior of reporters in cells in absence or presence of UV mutagenesis (right) to assess potential for His^+^ mutant generation on the His^-^ background, indicative of reporter suppression. **b.** Schematic of functional *hmo1Δ38::HIS3* reporter. *HMO1* promoter sequence between –405 and –39 was amplified and appended to the *HIS3* ORF followed by the *CYC1* terminator and integrated into a gene free region of chromosome I (chrI:199270-199271).

**Figure S2.**
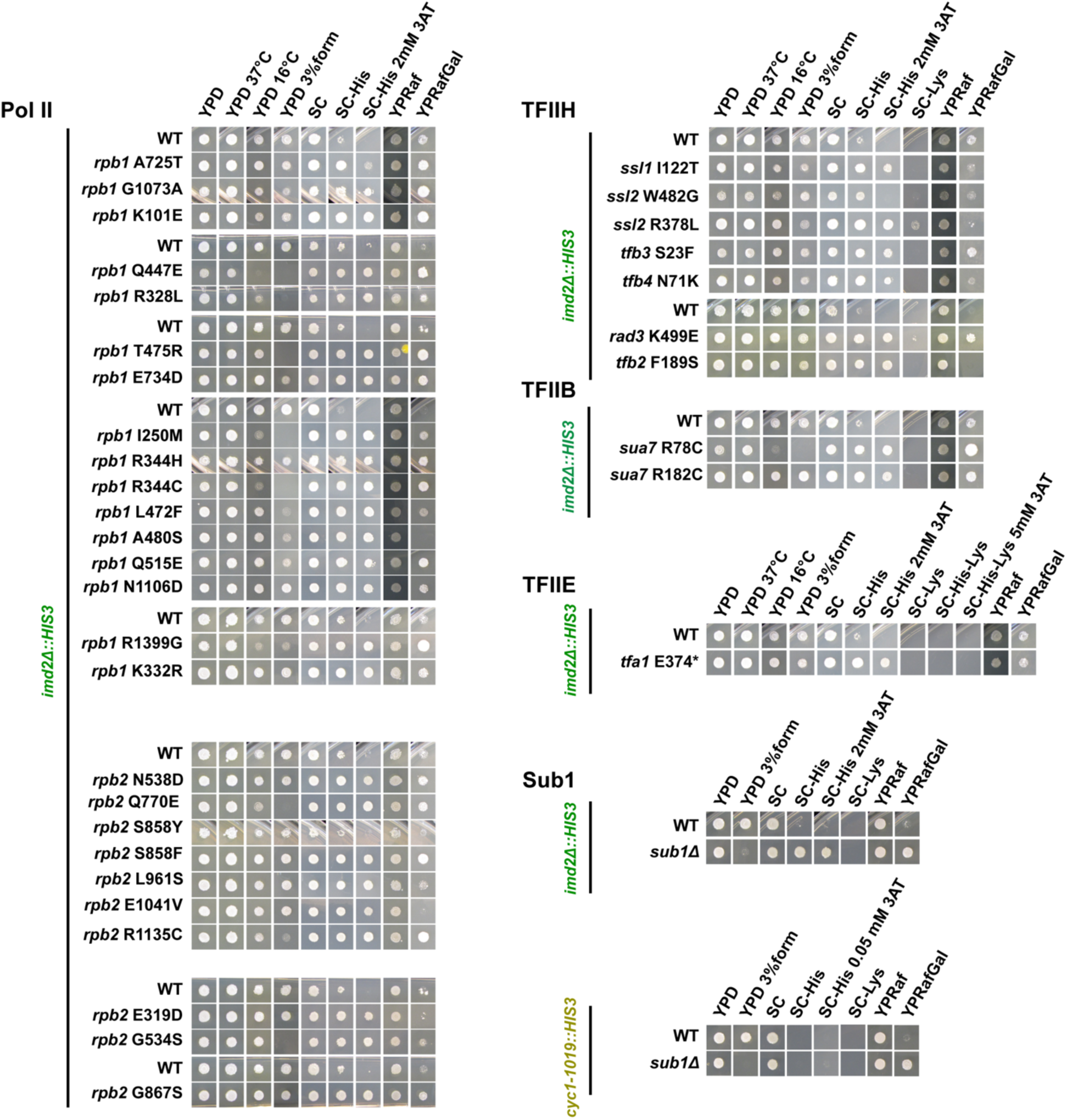
Initiation and secondary phenotypes of mutants isolated in genetic selections. Single spot phenotyping of a subset of downstream-shifting mutants isolated in our study.

**Figure S3.**
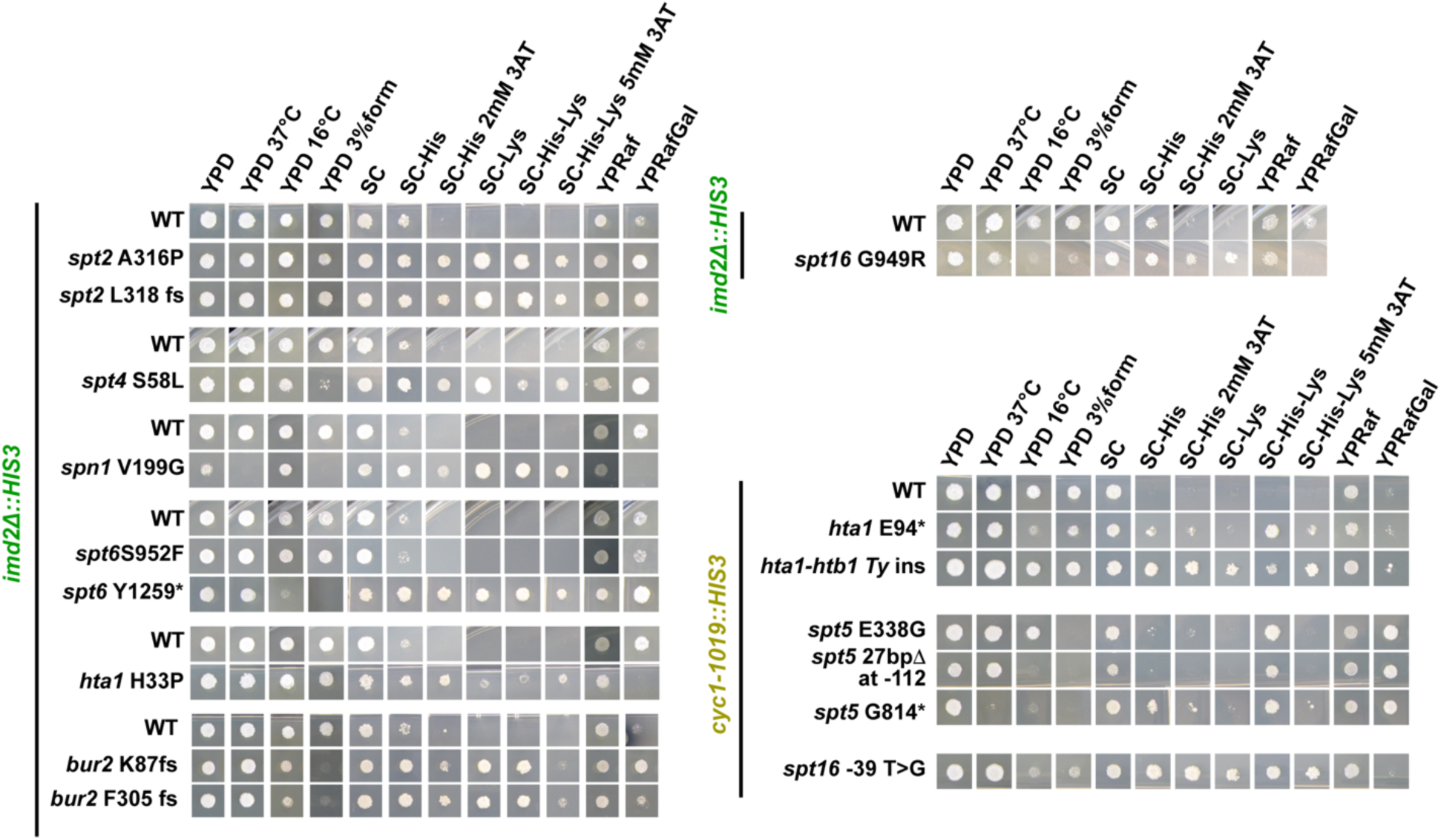
Phenotypes of chromatin mutants isolated in the study. Chromatin/Pol II elongation factor mutants identified for both downstream-shift sensitive reporters (subset shown).

**Figure S4.**
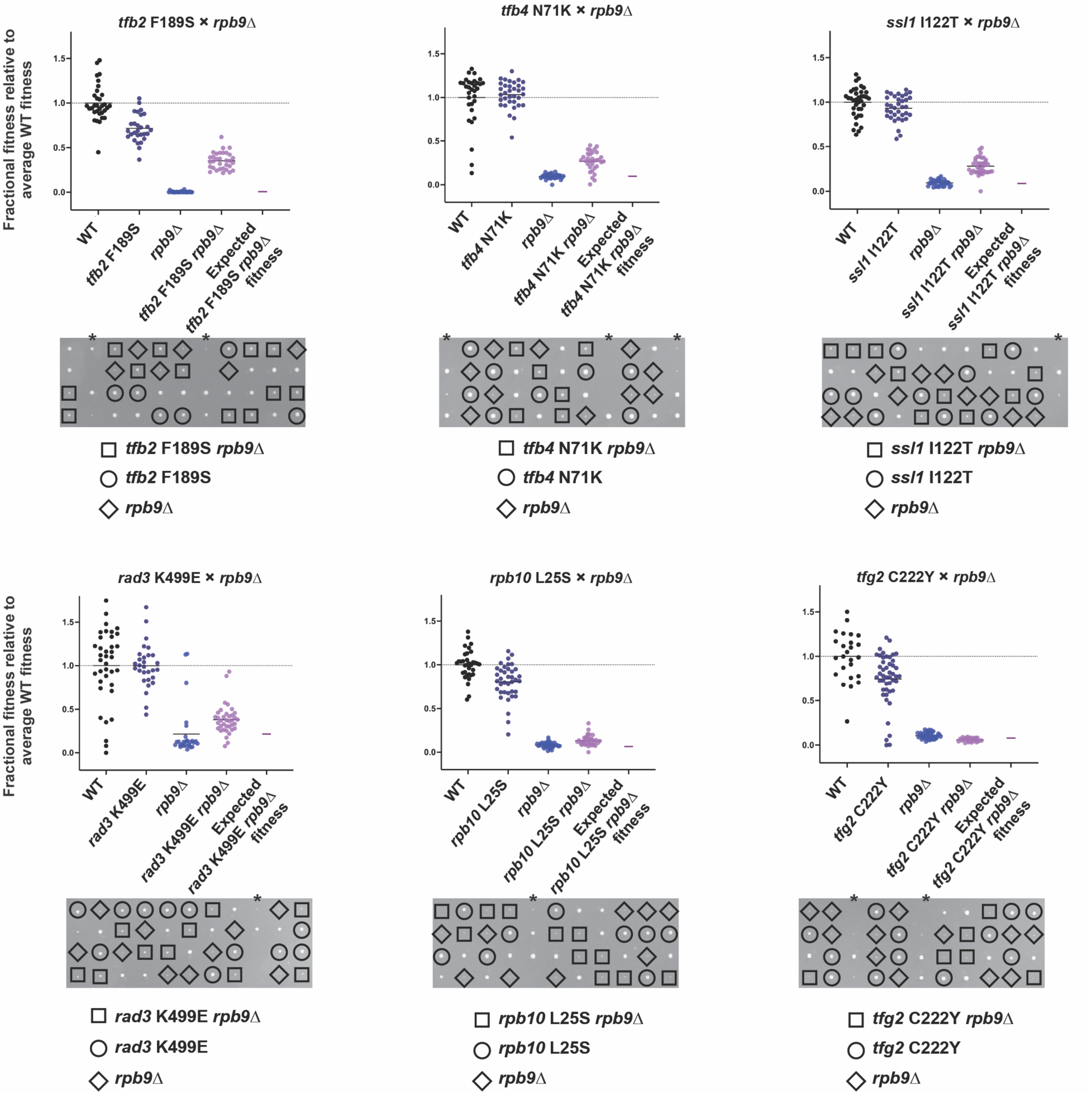
Genetic interactions between upstream-shifting *rpb9Δ* and novel downstream-shifting alleles prioritized in this study. Genetic interaction by tetrad analysis between *rpb9Δ* and downstream shifting novel alleles *tfb2* F189S, *tfb4* N71K, *ssl1* I122T across the top (left-right) and *rad3* K499E, *rpb10* L25S and *tfg2* C222Y across the bottom (left-right). Growth phenotypes quantified through spore colony size as in Figure 6. Suppression of *rpb9Δ* fitness defect by *tfb2* F189S, *tfb4* N71K, *ssl1* I122T and *rad3* K499E observed, while interactions with *rpb10* L25S and *tfg2* C222Y appear epistatic. Asterisks (*) indicate incomplete tetrads or failed dissections not considered.

**Figure S5.**
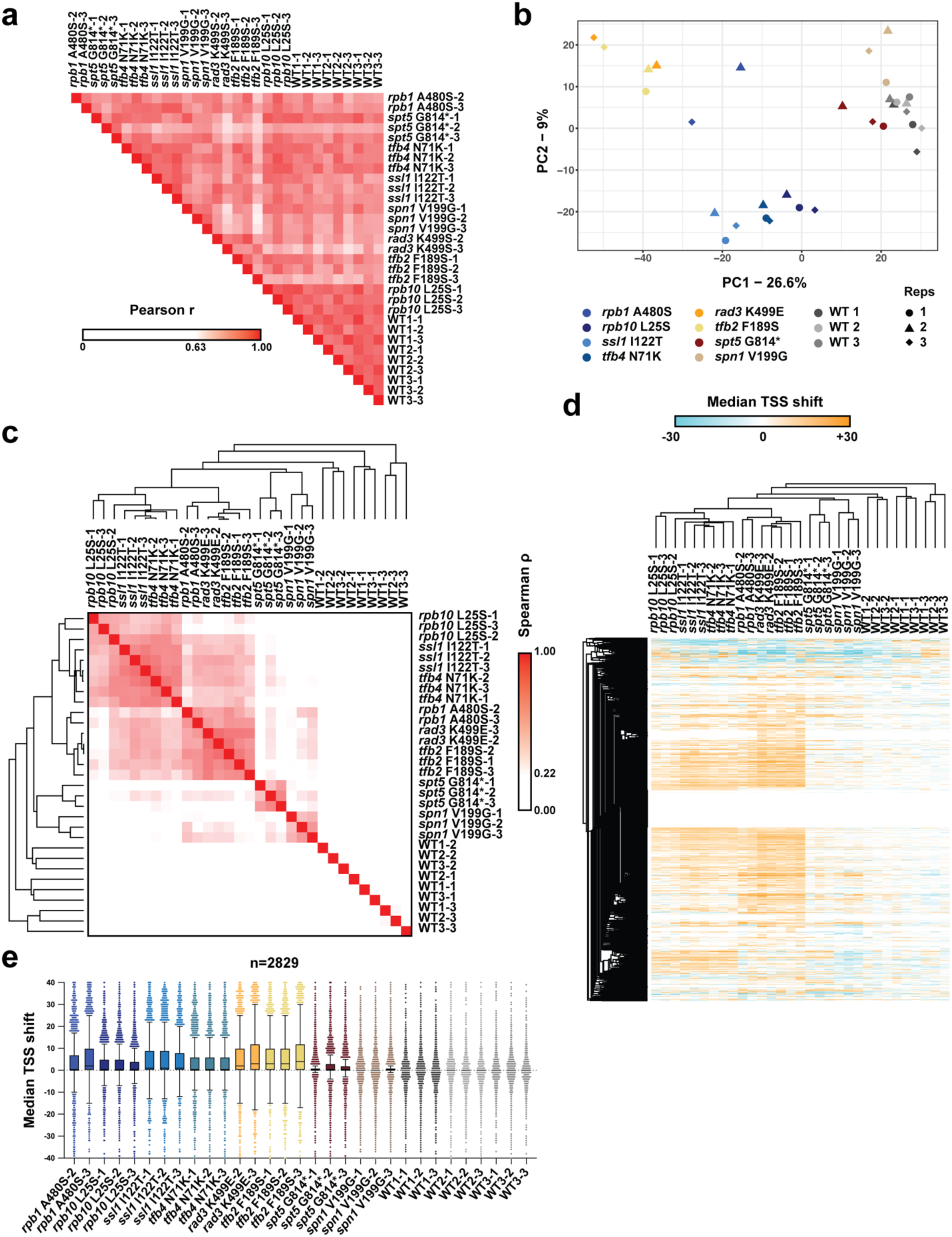
STRIPE-seq analysis of Δmedian values across individual mutant replicates. **a.** Pearson correlation of individual TSS positions between samples across genome with cutoff per position set at ≥ 3. **b.** Principal component analysis of Δmedian values across top 2829 genes qualifying expression cutoff of ≥10 total expression within 401 base analyzed promoter window for individual sample replicates. **c.** Spearman correlation of Δmedian of top 2829 genes across all samples and replicates. **d.** Hierarchically clustered heatmap of Δmedian values across all promoters in individually sequenced samples. **e.** Box-plot analysis of these Δmedian values amongst all sequenced samples analyzed by Tukey’s method.

**Figure S6.**
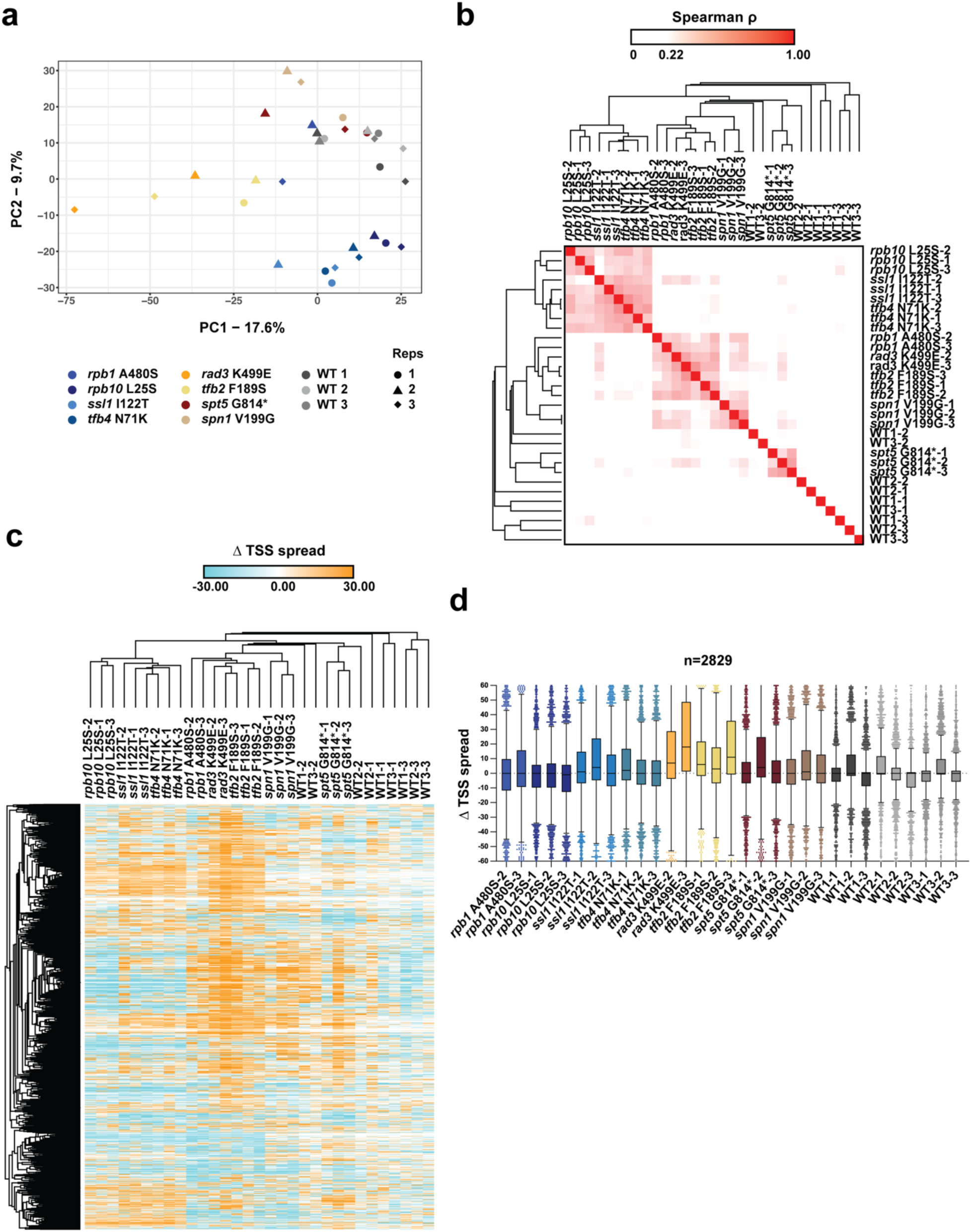
STRIPE-seq analysis of Δspread values across individual mutant replicates. **a.** Principal component analysis of Δspread values across top 2829 genes qualifying expression cutoff of ≥10 within 401 base analyzed promoter window for individual sample replicates. **b.** Pearson correlation of Δspread of top 2829 genes across all samples and replicates. **c.** Hierarchically clustered heatmap of Δspread values across all promoters in individually sequenced samples **d.** Box-plot analysis of these Δspread values amongst all sequenced samples analyzed by Tukey’s method.

## Notes

### Competing Interest Statement

The authors have declared no competing interest.

### Summary of Updates

Editing for clarity, additional genetic data added to Figure S4, Table S3 added

